# Lipid droplet accumulating microglia represent a dysfunctional and pro-inflammatory state in the aging brain

**DOI:** 10.1101/722827

**Authors:** Julia Marschallinger, Tal Iram, Macy Zardeneta, Song E. Lee, Benoit Lehallier, Michael S. Haney, John V. Pluvinage, Vidhu Mathur, Oliver Hahn, David W. Morgens, Justin Kim, Julia Tevini, Thomas K. Felder, Heimo Wolinski, Carolyn R. Bertozzi, Michael C. Bassik, Ludwig Aigner, Tony Wyss-Coray

**Affiliations:** Department of Neurology and Neurological Sciences, School of Medicine, Stanford University, Stanford, CA, USA; Paul F. Glenn Center for the Biology of Aging, Stanford University School of Medicine, Stanford, California, USA; Institute of Molecular Regenerative Medicine, Spinal Cord Injury and Tissue Regeneration Center Salzburg (SCI-TReCS), Paracelsus Medical University, Salzburg, Austria; Department of Genetics, School of Medicine, and Chemistry, Engineering, and Medicine for Human Health (ChEM-H), Stanford University, Stanford, CA, USA; Medical Scientist Training Program, Stanford University School of Medicine, Stanford, CA, USA; Department of Chemistry, Stanford University, Stanford, CA, USA; Department of Laboratory Medicine, Paracelsus Medical University, Salzburg, Austria; Obesity Research Unit, Paracelsus Medical University, Salzburg, Austria; Institute of Molecular Biosciences, BioTechMed-Graz, University of Graz, Graz, Austria; Stanford Neurosciences Institute, Stanford University, Stanford, CA, USA; Department of Veterans Affairs, Palo Alto, California, USA

## Abstract

Microglia become progressively activated and seemingly dysfunctional with age, and genetic studies have linked these cells to the pathogenesis of a growing number of neurodegenerative diseases. Here we report a striking buildup of lipid droplets in microglia with aging in mouse and human brains. These cells, which we call lipid droplet-accumulating microglia (LAM), are defective in phagocytosis, produce high levels of reactive oxygen species, and secrete pro-inflammatory cytokines. RNA sequencing analysis of LAM revealed a transcriptional profile driven by innate inflammation distinct from previously reported microglial states. An unbiased CRISPR-Cas9 screen identified genetic modifiers of lipid droplet formation; surprisingly, variants of several of these genes, including progranulin, are causes of autosomal dominant forms of human neurodegenerative diseases. We thus propose that LAM contribute to age-related and genetic forms of neurodegeneration.

## Introduction

Microglia are the resident immune cells of the central nervous system and play a pivotal role in the maintenance of brain homeostasis^1–3^. In the aging brain and in neurodegeneration, microglia lose their homeostatic molecular signature and show profound functional impairments, such as increased production of pro-inflammatory cytokines, elevated generation of reactive oxygen species (ROS) and build-up of dysfunctional lysosomal deposits indicative of impaired phagocytosis^4–9^. Recent single-cell transcriptome studies have revealed several distinct microglia subpopulations and cellular states in aging and disease^10–15^, including “disease associated microglia” (DAM)^12^, a presumably protective phagocytic microglia population, and “neurodegenerative microglia” (MGnD)^11^, a dysfunctional microglia phenotype. Furthermore, proliferative-region-associated microglia (PAM) arise during development and express genes that are also enriched in DAM^15^.

Over 100 years before these technologically-advanced genomic studies, Alois Alzheimer was likely one of the first to describe a unique microglial subset when he observed “many glial cells show[ing] adipose saccules” in brains of dementia patients (Alzheimer, 1907). Although microglia had not been identified as a distinct cell type back then, Alzheimer’s description of these cells suggests they were indeed microglia. Over the next few years, multiple studies confirmed this finding and regarded glial lipid accumulation as characteristic for senile dementia^16^. However, after this initial excitement, lipid deposits in microglia had mostly been ignored for almost a century.

Cellular lipid accumulation became of interest in other myeloid cells in the 1970s when “foamy macrophages” were discovered to contribute to atherosclerotic lesions^17, 18^. Since then, abnormal lipid accumulation has been recognized as a key aspect of immune dysfunction in myeloid cells^19^. In particular, lipid droplets, which are lipid storing organelles that contain neutral lipids such as glycerolipids and cholesterol, are increasingly accepted as structural markers of inflammation^19, 20^. Myeloid cells form lipid droplets in response to inflammation and stress, including the aforementioned macrophages in atherosclerotic lesions^21^, leukocytes in inflammatory arthritis^22^, and eosinophils in allergic inflammation^23^. Here, lipid droplets are production and storage sites for eicosanoids and inflammatory cytokines, and are further involved in antigen presentation and pathogen clearance^20, 24, 25^. Importantly, lipid-droplet rich foam cells in atherosclerosis show hallmarks of senescent cells and they seem deleterious at all stages of disease^26^.

Surprisingly, lipid droplets have not been studied functionally in brain myeloid cells in humans or vertebrates, and less than a handful of papers report the histological presence of lipid droplets in human brains^27–29^. That they may have important functions in disease has recently been suggested in a drosophila model where lipid droplet formation has been reported in glia at the onset of neurodegeneration^30, 31^. Oil-red-O positive “lipid-laden” cells, including neurons, astrocytes, ependymal cells and Iba1^+^ cells have recently been reported in mice and shown to increase with age ^32^. Lipid droplets were also induced in LPS-treated mouse hippocampal slice cultures and in the N9 microglia cell line^33–35^.

Together, lipid droplets have been recognized for their role as inflammatory organelles in peripheral myeloid cells, and lipid droplets in glia have been rediscovered in the context of brain aging and disease. Yet little is known about the formation and role of lipid droplets in microglia *in vivo*, and whether they have a role in neuroinflammation, brain aging, or neurodegeneration.

Here, we identify a novel state of microglia in the aging brain in which they accumulate lipid droplets. These lipid-droplet-accumulating microglia, which we named “LAM” in reference to recently described DAM – or disease-associated microglia^12^, exhibit a unique transcriptional signature, show defects in phagocytosis, produce increased levels of ROS and release elevated levels of pro-inflammatory cytokines. We identify *SLC33A1*, *SNX17*, *VPS35*, CLN, NPC2, and *GRN,* six genes with variants causing autosomal dominant forms of neurodegeneration, as genetic regulators of microglial lipid droplet formation, and we further validate the accumulation of LAM in GRN^-/-^ mice, a model for frontotemporal dementia. Together, these findings indicate that LAM represent a dysfunctional and pro-inflammatory microglia state in the aging brain, and further suggest that LAM might contribute to neurodegenerative diseases.

## Results

### Microglia accumulate lipid droplets in the aging brain

To determine whether microglia contain lipid droplets in the young and aged brain, we analyzed the cytoplasmic content of microglia from 3- and 20-month-old mice by transmission electron microscopy (EM). Interestingly, we frequently found lipid droplets (about 1 - 3 droplets per cell, often closely surrounded by lysosomes) in microglia of aged mice, but rarely detected them in young mice (Fig. 1a). For detailed quantitative analysis we co-localized TMEM119^+^ microglia with BODIPY, a dye that specifically labels neutral lipids and is commonly used to detect lipid droplets^36, 37^. We observed that in the aged brain, BODIPY^+^ microglia were abundant in the hippocampus but scarce in other regions (Supplementary Fig. 1), and therefore focused subsequent analyses on the hippocampus. Almost all lipid droplets in the hippocampus co-localized with TMEM119^+^ microglia, indicating that lipid droplets are primarily present in microglia but rare in other cell types. The percentage of BODIPY^+^ TMEM119^+^ microglia in the hippocampus was more than 4-fold higher in aged (51.95%) compared with young (12.08%) microglia, and lipid droplets were significantly larger in aged than in young microglia (Fig. 1b-e). To further validate the presence of lipid droplets, we used an antibody against the lipid droplet surface protein Perilipin 3 (Plin3) and detected Plin3^+^ droplets in aged TMEM119^+^ microglia (Fig. 1f). Next, we asked whether age-related microglial lipid droplet accumulation might also be relevant for human brain aging. We stained human postmortem hippocampus of young adult and aged cognitively normal individuals for Perilipin 2, a close paralog of Perilipin 3, and found that Plin2^+^ Iba1^+^ microglia were more frequent in aged than in young individuals (Fig. 1g).

**Fig. 1.**
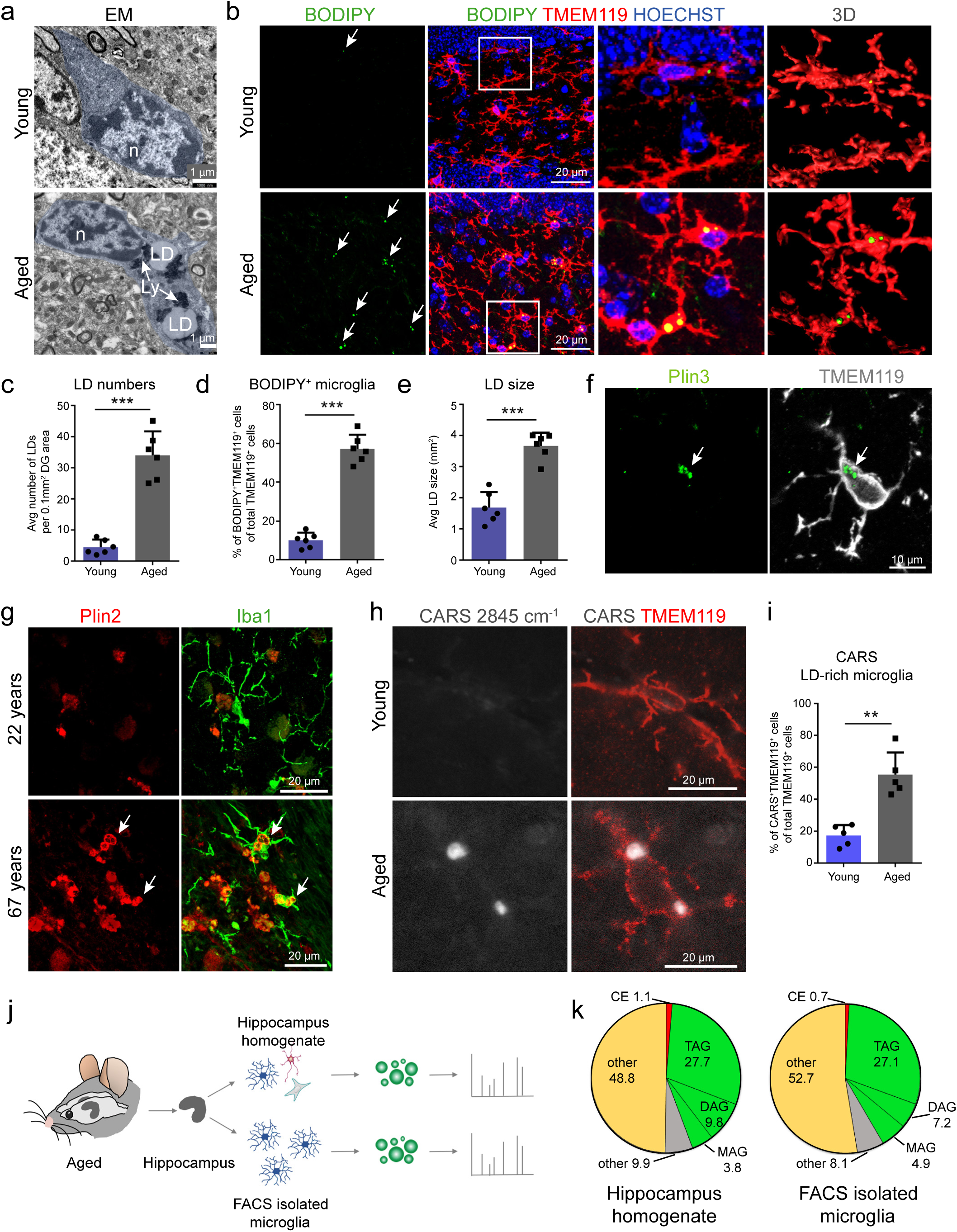
Microglia in the aged brain accumulate lipid droplets. **a**, Electron microscopy of microglia from young and aged mice. **b**, Hippocampus from aged mice stained for BODIPY^+^ (lipid droplets) and TMEM119^+^ (microglia). Right panel shows 3D reconstruction of BODIPY^+^/TMEM119^+^ microglia. Arrows indicate lipid droplets. **c-e**, Quantification of BODIPY^+^ lipid droplet numbers (**c**), percent BODIPY^+^/TMEM119^+^ cells (**d**), and average BODIPY^+^ lipid droplet size (**e**) in the hippocampus (dentate gyrus). n = 6 mice per group. **f**, Representative image of Plin3^+^ (lipid droplets) TMEM119^+^ microglia in aged mice. **g**, Confocal images of Plin2^+^ (lipid droplets) and Iba1^+^ (microglia) in the human hippocampus of a 22-year-old and 67-year-old individual. Arrows indicate Plin2^+^Iba1^+^ cells. **h**,**i**, Representative images (**h**) and quantification (**i**) of CARS^+^ signal (2845 cm^-1^) in TMEM119^+^ microglia in the hippocampus of young and aged mice. n = 5 mice per group. **j**,**k** Experimental schematic for lipidomics analysis of lipid droplets isolated from whole hippocampus and from FACS-sorted microglia from 20-month old mice (**j)**, and pie charts showing the composition of lipid droplets (**k**). n = 4 mice per group. Statistical tests: unpaired Student’s t-test. Error bars represent mean ± SD. **P< 0.01, ***P< 0.001. n, nucleus; LD, lipid droplet; Ly, Lysosome; TAG, triacylglycerol; DAG, diacylglycerol; MAG, monoacylglycerol; CE, cholesteryl ester. Young= 3-month-old male mice; Aged= 20 month-old male mice.

To corroborate our finding of lipid droplet accumulation in aged microglia, we used coherent anti-Stokes Raman scattering (CARS) microscopy, a label-free and nonlinear optical technique that enables the identification of types of molecules based on their specific vibrational energy. We performed CARS laser-scanning microscopy at 2,845 cm^-1^, which corresponds to the CH_2_ stretching frequency for neutral lipids and specifically identifies neutral lipids/lipid droplets^38^, on TMEM119^+^ immunostained brain sections from young and aged mice. Consistent with our previous data, we found that the numbers of CARS^+^ lipid storing microglia are significantly higher in aged than in young mice (50.76% vs 18.93%) (Fig. 1h,i).

Lipid droplets are composed of neutral lipids such as glycerolipids (triacylglycerols, diacylglycerols, monoacylglycerols) and cholesteryl esters, yet their content can vary greatly between cell types. We isolated lipid droplets from whole hippocampi of aged mice and, to more specifically determine the content of microglial lipid droplets, from FACS-sorted aged hippocampal microglia (Fig. 1j). Lipidomics analysis revealed that lipid droplets from the whole hippocampus and from aged microglia show a nearly identical lipid distribution and are mainly composed of glycerolipids (hippocampus: 41.3%; microglia: 44.4%), while cholesteryl esters were almost absent (hippocampus: 1.1%; microglia: 0.7%) (Fig. 1k).

### Lipid droplet-rich microglia have a unique transcriptome signature that is associated with cellular dysfunctions

To determine the transcriptional phenotype of lipid droplet-containing microglia in the aged brain, we isolated CD11b^+^CD45^lo^ microglia from the hippocampi of 18-month old mice based on their BODIPY^+^ mean fluorescence intensity and analyzed lipid droplet-low (BODIPY^lo^; LD-lo) and lipid droplet-rich (BODIPY^hi^; LD-hi) microglia by RNA-Sequencing (RNA-Seq) (Fig. 2a,b). Of note, we used an optimized microglia isolation strategy that uses mechanical tissue homogenization instead of enzymatic digestion, which keeps microglia largely in a non-activated state and therefore prevents unwanted bias towards an activated pro-inflammatory signature^39^. Unsupervised cluster analysis segregated LD-lo from LD-hi microglia and revealed prominent differences between their transcriptome with 692 significantly differentially expressed genes (Fig. 2c,d).

**Fig. 2.**
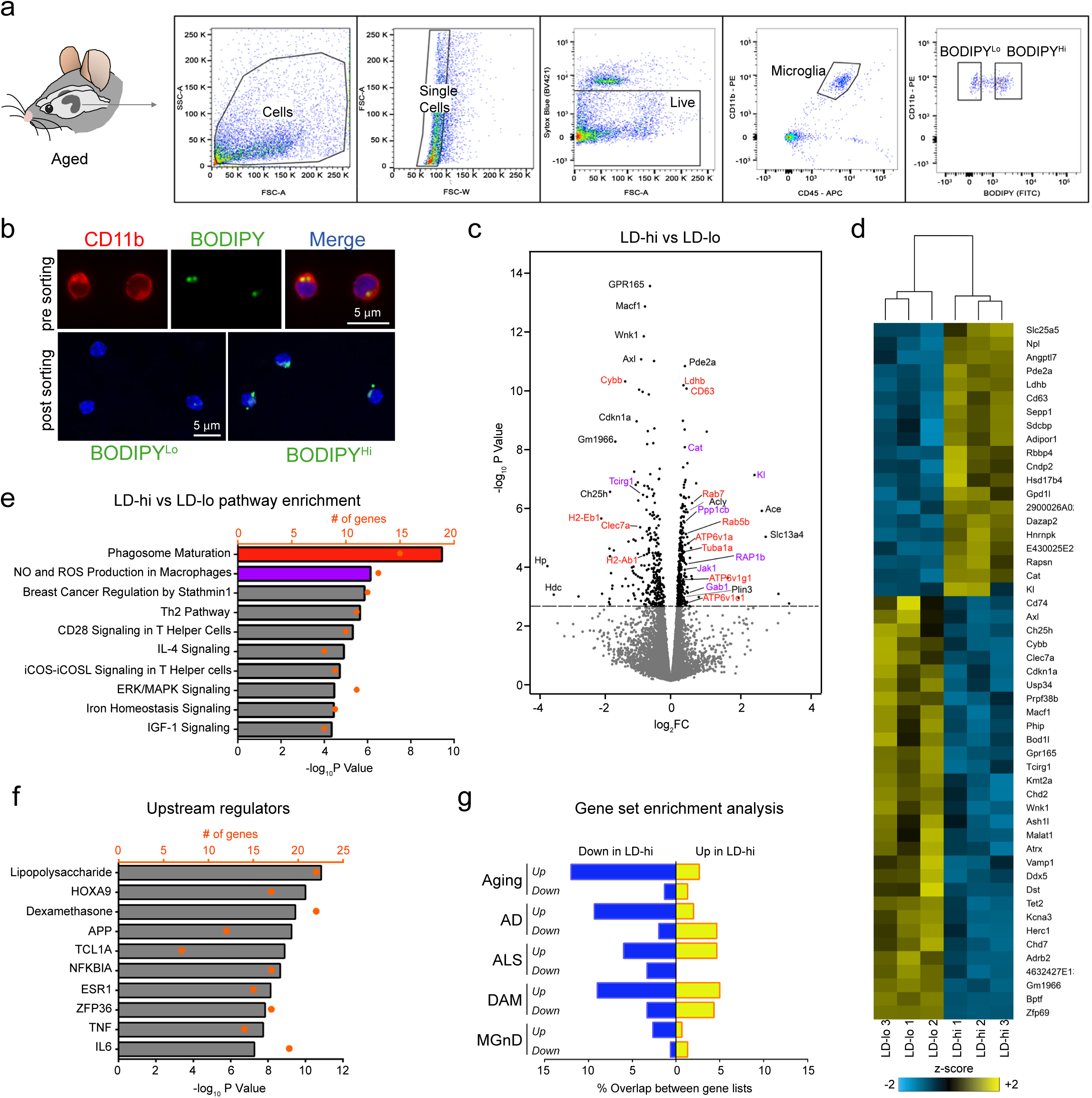
RNA-Seq of lipid droplet-low and lipid droplet-high microglia from aged mice reveals transcriptional changes linked to phagocytosis and ROS production. **a**, Flow sorting scheme for isolation of BODIPY^lo^ (=LD-lo) and BODIPY^hi^ (=LD-hi) CD11b^+^CD45^lo^ cells from the hippocampus of 18-month old male mice. n = 3 samples per group. Each sample is a pool of microglia from 3 mouse brains. **b**, Representative images of microglia after brain homogenization and marker staining, before (upper panel) and after (lower panel) FACS sorting. **c**, Volcano plot showing differentially expressed genes in LD-hi versus LD-lo microglia. Dashed line represents q-value < 0.05 cutoff. Genes involved in phagosome maturation (red) and ROS production (purple) are highlighted. **d**, Heatmap showing the top 50 differentially expressed genes (ranked by p-value). **e**, Top canonical pathways identified by IPA that are differentially regulated between LD-hi and LD-lo microglia. Analysis based on top 200 genes ranked by p-value. **f**, IPA upstream regulator analysis of top 200 differentially expressed genes between LD-lo and LD-hi microglia. **g**, Overlap between genes changing in microglia in aging and neurodegeneration (Aging, AD, ALS, DAM, MGnD), and genes upregulated (yellow) or downregulated (blue) in LD-hi microglia. Percent overlap denotes the fraction of genes in each gene list that are up- or down-regulated in LD-hi microglia. LD, lipid droplet.

Pathway Analysis of differentially expressed genes revealed *Phagosome maturation* and *Production of NO and ROS*, two key functions of microglia that become dysregulated with age, as the most significant pathways associated with LD-hi microglia (Fig. 2e). Regulated genes in the phagosome maturation pathway included lysosomal genes (*CD63*, *ATP6V1A*, *ATP6V1C1*, *ATP6V1G1*, *TUBA1*), genes involved in vesicular transport (*RAB5B*, *RAB7*), and *CD22*, a negative regulator of phagocytosis in microglia^80^. Interestingly, most genes linked to NO and ROS generation (e.g. *CAT*, *KL*, *PPP1CB*, *JAK*, *RAP1B*) were upregulated in LD-hi microglia (Fig. 2c). In addition, LD-hi microglia were enriched in lipid related genes, including the lipid droplet specific gene Perilipin 3 (*PLIN3*) and the ATP citrate synthase *ACLY*, which is involved in lipogenesis (Supplementary Table 1), and *fatty acid* β*-oxidation* was one of the top enriched pathways related to genes upregulated in LD-hi microglia (Supplementary Fig. 2). Next, we used annotated functional transcriptomics to predict upstream activators of genes differentially regulated between LD-lo and LD-hi microglia. The pro-inflammatory endotoxin lipopolysaccharide (LPS) was the most significant upstream regulator, suggesting a link between innate inflammation and lipid droplets in microglia (Fig. 2f).

Lastly, we compared the transcriptional profile of LD-hi microglia with that of microglia in aging^40^, ALS^41^, and AD^42^, and of microglia subpopulations recently identified in development, aging and disease, including “disease-associated microglia” (DAM)^12^, “neurodegenerative microglia” (MGnD)^11^, and microglia clusters revealed by Li et al. (2019) and Hammond et al. (2019)^14, 15^. We found a moderate overlap between genes differentially expressed in LD-hi microglia and the Cluster 3 microglia from Hammond and colleagues (2019)^14^, a microglia subset mainly detected in E14.5 brains with a transcriptional signature linked to inflammation and metabolic pathways (Supplementary Figure 2c-i). Furthermore, genes downregulated in LD-hi microglia overlapped partially with published gene sets of microglia in aging and neurodegeneration. (Fig. 2g). However, the downregulated genes of LD-hi microglia matched primarily with genes that were upregulated in microglia in aging and in DAM (e.g. *AXL*, *CD74*, *CLEC7A*, *CYBB*) (Supplementary Fig. 2).

Overall, these data suggest that lipid droplet-containing aged microglia show transcriptional changes of genes related to key microglia functions such as phagocytosis, ROS production and immune signaling, yet they have a unique transcriptome signature that is distinct from previously described microglia states observed in aging and neurodegeneration. We therefore designate this microglia state as “lipid droplet accumulating microglia” (LAM) and will use this term henceforth.

### The innate TLR4 ligand LPS induces lipid droplet formation in microglia

LPS is the main upstream regulator of genes differentially expressed between LD-lo and LD-hi microglia (Fig. 2f), and immune cells such as macrophages, neutrophils, and eosinophils accumulate lipid droplets in response to inflammatory conditions^20^. To determine whether inflammation triggers lipid droplet formation in microglia, we treated the mouse microglia-derived cell line BV2 with LPS and found a 5-fold increase of BODIPY^+^ BV2 cells compared with control cells (Fig. 3a,b). To confirm the identity of these BODIPY-labeled droplets we used Triacsin C, an inhibitor of long-chain acyl-CoA synthetase which inhibits *de novo* synthesis of glycerolipids and prevents lipid droplet formation^35, 47^. Indeed, treatment with Triacsin C abolished the LPS-induced increase in BODIPY^+^ cells (Fig. 3a,b). Accordingly, LPS-treated BV2 cells showed significantly increased BODIPY^+^ fluorescence when analyzed by flow cytometry (Fig. 3c,d), and co-treatment with Triacsin C inhibited an increase in BODIPY^+^ signal in LPS-stimulated BV2 cells (Fig. 3c,d).

**Fig. 3.**
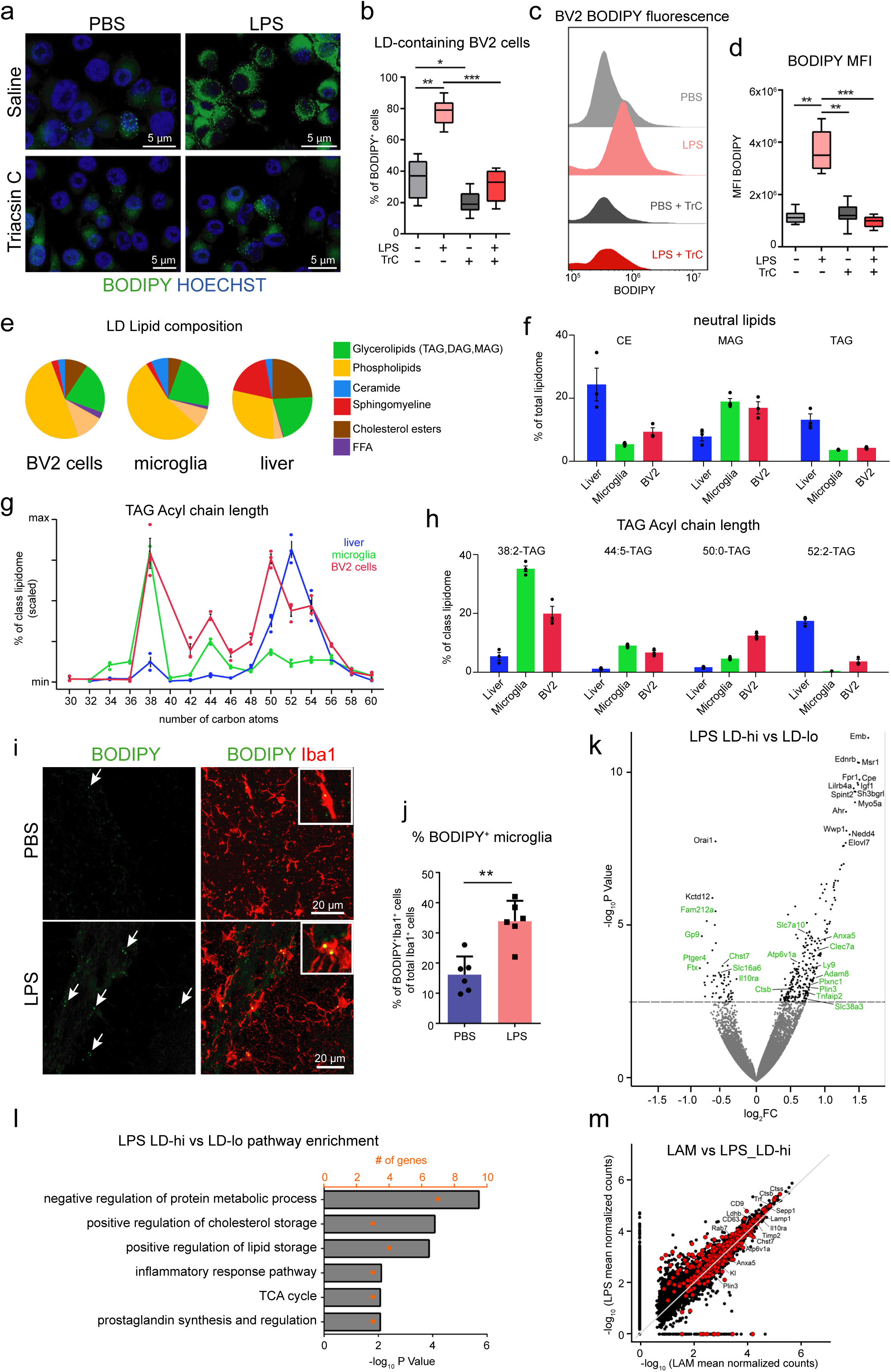
LPS treatment induces lipid droplet formation in microglia. **a-d**, BV2 cells were treated with PBS or LPS (5 μg/ml) for 18 hours and co-treated with Triacsin C (1μM) or saline. Representative micrographs of BODIPY staining in BV2 cells (**a**) and quantification of BODIPY^+^ cells (**b**). **c**,**d**, Representative flow cytometry histogram (**c**) and quantification (**d**) of BODIPY fluorescence in BV2 cells. **e**-**h**, Lipidome profiling of lipid droplets from LPS-treated BV2 cells, and from primary microglia and liver tissue from aged (20-month-old) mice. Overall composition of lipid droplets (**e**), percentage of neutral lipids (**f**), and distribution of TAG species (**g**,**h**). **i**,**j**, BODIPY^+^ and Iba1^+^ in the hippocampus of 3-month old male mice given intraperitoneal (i.p.) injections of PBS or LPS (1 mg/kg BW) for four days. Representative confocal images (**i**) and quantification (**j**) of BODIPY^+^/Iba1^+^ microglia. n = 6 mice per group. **k**-**m**, RNA-Sequencing of BODIPY^lo^ (=LD-lo) and BODIPY^hi^ (=LD-hi) CD11b^+^CD45^lo^ microglia from the hippocampus of 3-month old LPS-treated mice. n = 4 samples per group. Each sample is a pool of microglia from 2 mouse brains. **k**, Volcano plot showing differentially expressed genes in LD-hi versus LD-lo microglia. Dashed line represents q-value < 0.05 cutoff. LAM genes are highlighted in green. **l**, EnrichR pathway analysis of genes differentially regulated between LD-hi and LD-lo microglia. **m**, Scatterplot showing gene expression intensities (mean normalized counts) of LD-hi microglia in LPS-treated mice compared to LAM in aging. Genes differentially expressed in LAM are highlighted in red. Experiments on BV2 cells were performed three times in technical triplicates. Statistical tests: unpaired Student’s t-test (j) and one-way ANOVA followed by Tukey’s post hoc test (b,d). Error bars represent mean ± SD (j). Horizontal lines in the box plots indicate medians, box limits indicate first and third quantiles, and vertical whisker lines indicate minimum and maximum values (b,d). *P< 0.05, **P< 0.01, ***P< 0.001. CE, cholesteryl ester; DAG, diacylglycerol; LD, lipid droplet; MAG, monoacylglycerol; MFI, mean fluorescent intensity; TAG, triacylglycerol; TrC Triacsin C.

To explore whether LPS-induced lipid droplets in BV2 cells resemble those in microglia from aged mice, we performed lipidomics of lipid droplets isolated from BV2 cells and from aged microglia. Interestingly, they had a highly similar overall lipid content and a similar composition of neutral lipids, with low amounts of cholesteryl esters but high levels of glycerolipids (MAG, TAG). In contrast, lipid droplets from the liver, which we included as a control because this tissue shows a strong accumulation of lipids, contained high levels of cholesteryl esters (Fig. 3e,f, Supplementary Fig. 3). In addition, lipid droplets from BV2 cells and from aged microglia showed a similar chain length distribution of TAG-associated fatty acids with peaks at 38:2 and 44:5, while liver lipid droplets had a shift towards longer TAG chain fatty acids with a peak at 52:2 carbon atoms, which is in agreement with previous reports^81^ (Fig. 3g,h).

We next treated young mice, which contain low numbers of lipid droplet-containing microglia (Fig. 1b), systemically with LPS at a dose that has been previously shown to induce a pro-inflammatory phenotype in microglia (1 mg LPS/kg BW for four days^46^). Notably, we observed a significant, twofold increase in lipid droplet-containing microglia (BODIPY^+^Iba1^+^) in the hippocampus of LPS-treated mice compared with vehicle treated controls (Fig. 3i,j). To test whether LD-hi microglia from LPS-treated mice were transcriptionally similar to LAM, we performed RNA Sequencing on BODIPY^lo^ (LD-lo) and BODIPY^hi^ (LD-hi) microglia from LPS-treated young mice. Unsupervised cluster analysis revealed 272 differentially expressed genes (Fig. 3k), and enriched pathways were linked to metabolism, lipid storage, and inflammation (Fig. 3l). We compared gene expression of LD-hi microglia from LPS-treated mice and of LAM in aging, and found that expression intensities for genes differentially expressed in LAM were highly similar in both datasets (Fig 3m).

Together, these findings demonstrate that LPS triggers lipid droplet formation in microglia *in vitro* and *in vivo*, and lipid droplet containing microglia in young LPS-treated mice show a transcriptional signature that partially overlaps with LAM. Furthermore, LPS-induced lipid droplets in BV2 cells are highly similar to lipid droplets in aged microglia, thus making this *in vitro* assay a useful model to study lipid droplets in LAM.

### LAM have phagocytosis deficits

Impaired phagocytosis is considered a hallmark of microglia aging^48^. Because *Phagosome maturation* was the top regulated pathway associated with the transcriptome of LAM (Fig. 2e), we sought to examine whether the phagocytosis pathway was altered in these cells. Our RNA-Seq analysis revealed an upregulation of endosomal and lysosomal genes (Fig. 4a), suggesting that LAM show an increased load of endosomal/lysosomal vesicles. To corroborate this hypothesis, we analyzed immunoreactivity for the lysosome-associated protein CD68 in in the hippocampus from aged mice and found that BODIPY^+^ microglia had significantly higher CD68 expression than BODIPY^-^ microglia (Fig. 4b,c). Furthermore, 3D reconstruction revealed that CD68^+^ vesicles often accumulate closely around lipid droplets (Fig. 4d). Electron microscopy confirmed an accumulation of lysosomes in lipid droplet containing microglia (Fig. 4e).

**Fig. 4.**
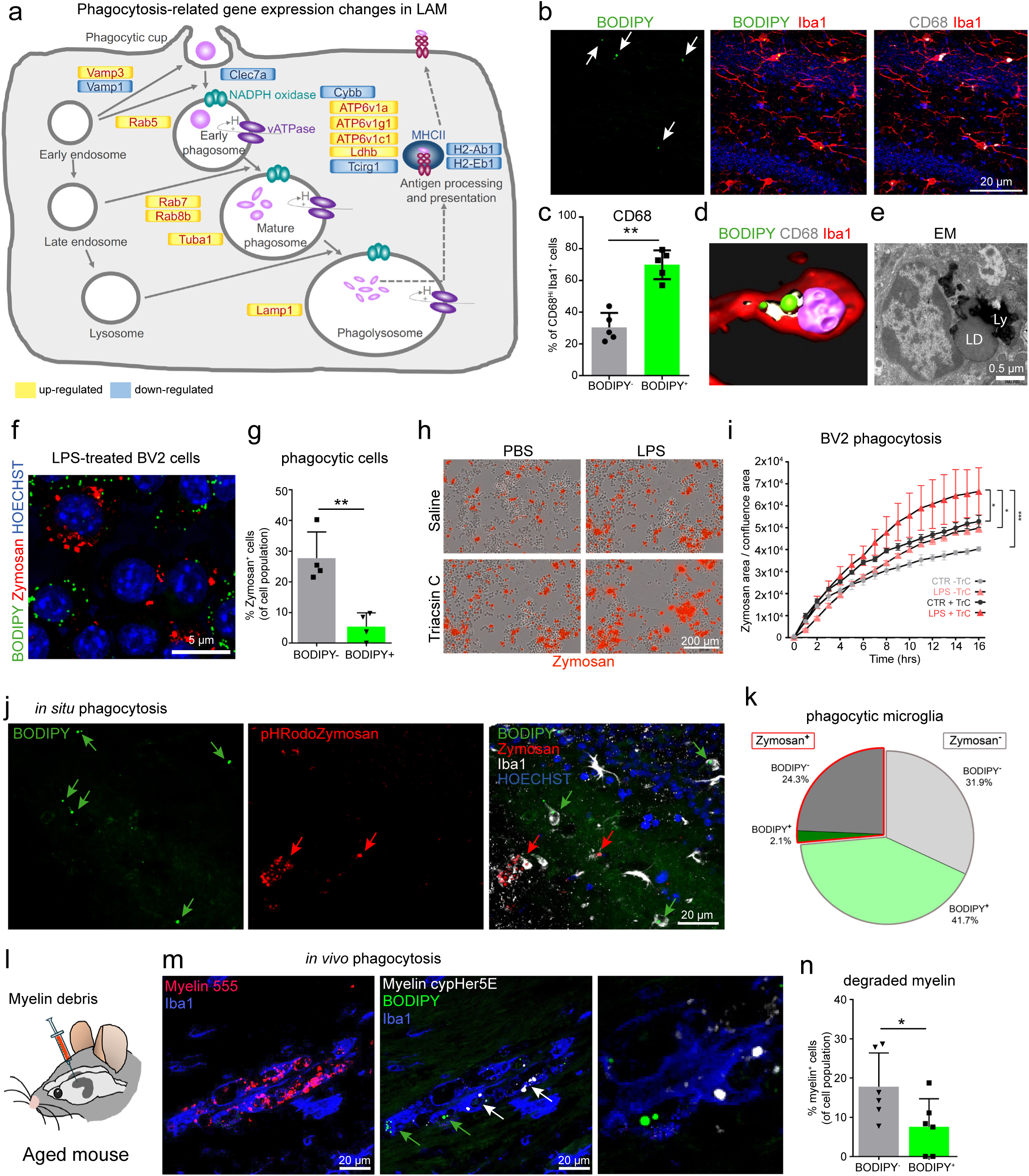
LAM and lipid droplets in BV2 cells are associated with impaired phagocytosis. **a**, Pathway map of genes related to phagosome maturation that are differentially expressed in LAM (see Fig. 2). **b**, Confocal images showing BODIPY^+^ (lipid droplets), CD68^+^ (endosomes/ lysosomes), and Iba1^+^ in the hippocampus from 20-month old mice. **c**, Percentage of BODIPY^-^ and BODIPY^+^ Iba1^+^ microglia with high levels of CD68 (CD68^hi^). n = 5 mice per group. **d**, 3D reconstruction of Iba1^+^ microglia showing BODIPY^+^ lipid droplets closely surrounded by CD68^+^ vesicles. **e**, Electron microscopy image showing lysosomal accumulation in LAM from a 20-month old mouse. **f**,**g**, Confocal images (**f**) and quantification (**g**) of BODIPY^+^ and Zymosan^+^ in BV2 cells treated with LPS (5 ug/ml) for 18 hours. **h**,**i**, Phagocytosis of pHrodo red Zymosan in BV2 cells treated with PBS or LPS and co-treated with Triacsin C (1μM) or saline. Representative images of (**h**) and time lapse imaging and quantification (**i**) of Zymosan uptake in BV2 cells. **j,k**, 250 μm organotypic brain slices from 12-month old mice were incubated for 4 hours with pHrodo red Zymosan particles. Representative confocal images of the hippocampus (**j**) and pie chart showing the percentages of Zymosan-containing BODIPY^-^ and BODIPY^+^ Iba1^+^ cells (**k**). P-value for Zymosan^+^BODIPY^-^ vs Zymosan^+^BODIPY^+^ Iba1^+^ cells = 0.0012. n = 3 mice per group. **l-n**, Myelin debris co-labelled with Alexa Fluor 555 and with CypHer5E pH-dependent fluorescent dye was stereotactically injected into the hippocampus of 20-month old male mice, and phagocytosis was analyzed 48 hours after injection. **m**, Representative images of AF555-labelled myelin (left panel) and of Iba1^+^ cells with and without lipid droplets (BODIPY^+^) and acidified myelin (Myelin cypHer5E^+^). **n**, Quantification of myelin uptake in BODIPY^+^/Iba1^+^ and BODIPY^-^ Iba1^+^ cells. n = 6 mice. Experiments on BV2 cells were performed three times in technical triplicates. Statistical tests: unpaired Student’s t-test (c,g,n), two-way ANOVA followed by Tukey’s post hoc test (i,k). Error bars represent mean ± SD (c,g,n) and mean ± SEM (i). *P< 0.05, **P< 0.01, ***P< 0.001. LD, lipid droplet; Ly, Lysosomes; n, nucleus; TrC, Triacsin C.

To determine phagocytic activity in lipid droplet-containing microglia and whether modulation of lipid droplets affects this function, we used the BV2 *in vitro* model described earlier (Fig. 3a,b) to induce (LPS) or to inhibit (Triacsin C) lipid droplet formation, and analyzed the pHrodo red Zymosan uptake in these cells (Fig. 4f-i). In line with previous reports^49^, LPS increased phagocytosis (Fig. 4h,i) yet, interestingly, Zymosan particles were mainly found in the BODIPY^-^ cell population and to a significantly lesser extent in lipid droplet-rich BODIPY^+^ cells (Fig. 4f,g). Furthermore, Triacsin C increased Zymosan phagocytosis in LPS-treated cells (Fig. 4h,i). Of note, to test lipid droplet formation in BV2 cells and its effects on phagocytosis in a system that mimics the aging environment, we treated BV2 cells with aged plasma, which has been shown to activate microglia and to trigger brain aging^82^. Indeed, aged plasma induced lipid droplet formation, and again, cells with high numbers of lipid droplets showed significantly less Zymosan uptake (Supplementary Fig. 4).

Next, to analyze phagocytosis in LAM, we prepared acute organotypic brain slices from 12-month-old mice and treated them with pHrodo red Zymosan particles. pHrodo Zymosan becomes fluorescent at an acidic pH and thus serves as a sensor for lysosomal degradation of phagocytosed material. Remarkably, the percentage of Zymosan^+^ BODIPY^+^ microglia was ten-fold lower compared to Zymosan^+^ BODIPY^-^ microglia (2.1% vs 24.3%), indicating that LAM have severe defects in degrading Zymosan (Fig. 4j,k). In addition, we assessed phagocytic activity of LAM *in vivo* by injecting myelin debris, a phagocytic substrate that accumulates in the aging brain, into the hippocampus of aged mice (Fig 4l). We labelled the myelin debris with a constitutively fluorescent dye (A555) and with a pH-sensitive dye (cypHer5E) to detect its lysosomal degradation, and found that LAM phagocytosed significantly fewer cypHer5E^+^ myelin particles compared to microglia without lipid droplets (Fig. 4m,n).

Collectively, these data suggest that lipid droplet-rich microglia have phagocytosis deficits and that increased lipid storage is associated with impaired phagocytosis.

### LAM produce high levels of ROS and show excessive release of pro-inflammatory cytokines

Aged microglia are one of the main sources of increased ROS levels in the aging brain ^8, 48, 50^, and excessive microglial ROS production might contribute to age-related CNS dysfunctions. Intriguingly, pathway analysis of genes differentially expressed between aged LD-lo and LD-hi microglia identified *Production of NO and ROS* as the second most significantly regulated pathway (Fig. 2e). Given that over 90% of the differentially expressed genes related to ROS and NO production were upregulated (Fig. 5a), we hypothesized that LAM generate increased levels of ROS.

**Fig. 5.**
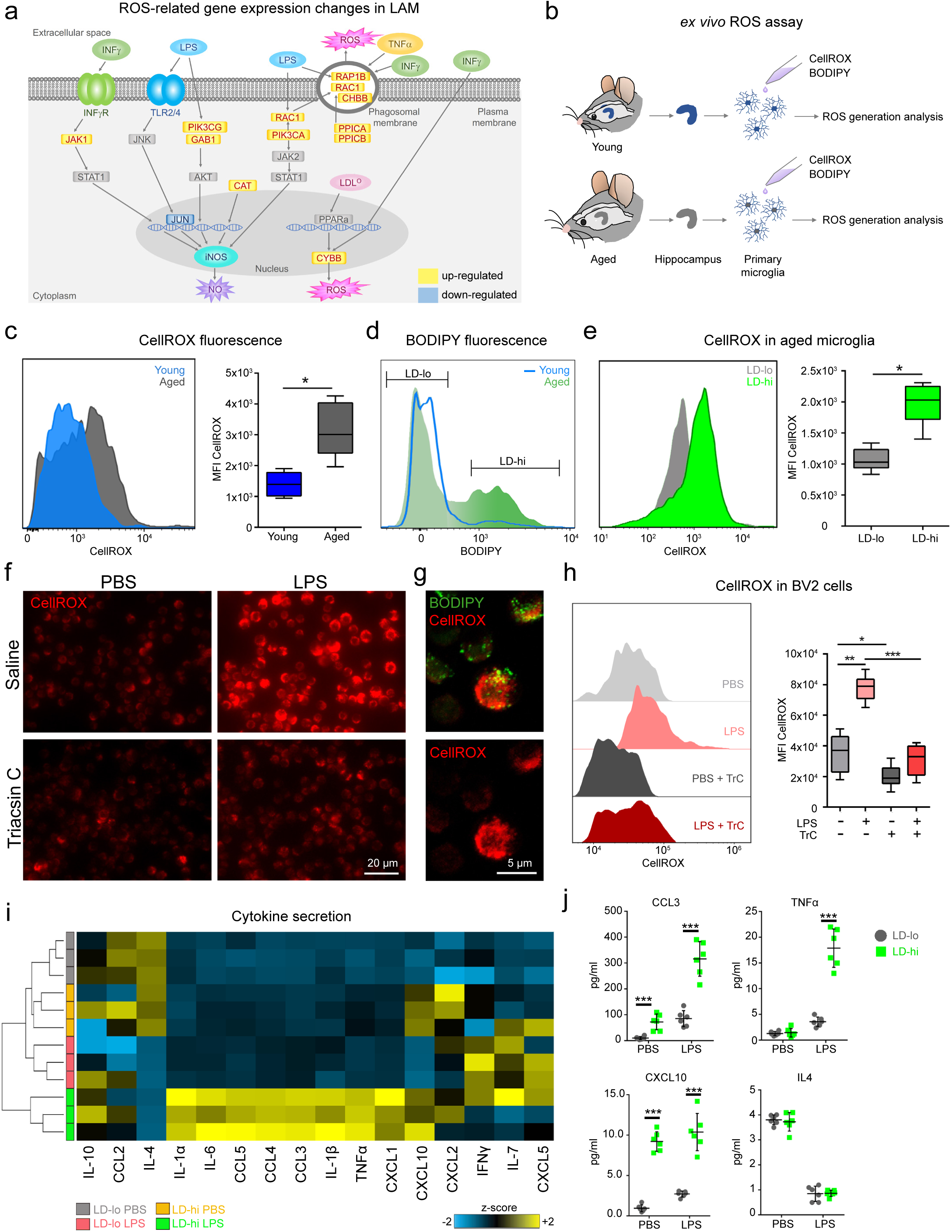
LAM and lipid droplet-rich BV2 cells show increased ROS production and LAM secrete elevated levels of inflammatory cytokines. **a**, Pathway map of genes related to ROS production that are differentially expressed in LAM (see Fig. 2). **b**, Experimental schematic of ROS analysis in primary microglia from young and aged mice. **c**, Representative flow cytometry histogram and quantification of CellROX fluorescence in primary microglia from young and aged mice. **d**, Gating scheme for BODIPY^lo^ (LD-lo) and BODIPY^hi^ (LD-hi) microglia from aged mice. **e**, Histogram and quantification of CellROX fluorescence in LD-lo and LD-hi microglia from aged mice. **f**-**h**, CellROX fluorescence in BV2 cells treated with PBS or LPS (5 ug/ml) for 18 hours, co-treated with Triacsin C (1μM) or saline. **f**, Representative images of CellROX^+^ signal in BV2 cells. **g**, Confocal images showing BODIPY^+^ and CellROX^+^ in LPS treated BV2 cells. **h**, Flow cytometry histogram and quantification of CellROX fluorescence in BV2 cells. **i,j** Acutely isolated LD-lo and LD-hi primary microglia from aged mice were treated with LPS (100 ng/ml) for 8 hours, and cytokine concentrations in the medium were measured using multiplex array. Heatmap showing changes in cytokine secretion under baseline conditions and after LPS treatment (**i**) and individual dot plots of selected cytokines (**j**). Experiments were performed three (BV2 cells) or two (primary cells) times in technical triplicates. Primary cells were isolated from three mice per group per experiment. Statistical tests: unpaired Student’s t-test (c,e), one-way ANOVA (h,j) followed by Tukey’s post hoc test. Horizontal lines in the box plots indicate medians, box limits indicate first and third quantiles, and vertical whisker lines indicate minimum and maximum values. *P< 0.05, **P< 0.01, ***P< 0.001. LD, lipid droplet; MFI, mean fluorescent intensity; TrC, Triacsin C. Young= 3-month-old male mice; Aged= 20 month-old male mice.

First, we sought to test whether ROS production was changed in aged compared to young microglia in general. We stained single cell suspensions from the hippocampi of young and aged mice with CellROX, a dye that is non-fluorescent in a reduced state but exhibits bright fluorescence upon oxidation by ROS, and analyzed CellROX fluorescence within the CD11b^+^CD45^lo^ microglia population by flow cytometry (Fig. 5b). In line with previous reports^51^, we found that aged microglia had significantly higher ROS levels than young microglia (Fig. 5c).

Next, we measured ROS levels in LD-lo and LD-hi microglia from aged mice and found that CellROX signal was twofold increased in LD-hi compared to LD-lo microglia (Fig. 5d-e), suggesting that the elevated ROS levels in aged microglia might be specifically driven by the increased ROS generation of LAM.

To examine whether modulation of lipid droplet numbers would affect ROS production, we used LPS to induce lipid droplets in BV2 cells and inhibited lipid droplet formation by co-treatment with Triacsin C (Fig. 3a,b). We assessed CellROX fluorescence using microscopy and flow cytometry. In line with previous reports, LPS treatment induced elevated ROS generation in BV2 cells^52^. Interestingly, inhibition of lipid droplets with Triacsin C was sufficient to significantly reduce ROS levels in LPS-treated cells (Fig. 5f,h). Moreover, high magnification confocal imaging of LPS-treated BV2 cells revealed that cells with high CellROX fluorescence were often loaded with lipid droplets (Fig. 5g). Together, these findings demonstrate that LAM have elevated concentrations of ROS and suggest that lipid droplets mediate LPS-induced ROS generation.

Another characteristic of aged microglia is the increased production of inflammatory cytokines such as TNF-α, IL-1β and IL-6 under baseline conditions and excessive cytokine release upon immune challenge^48^. To determine the cytokine expression profile of LAM, we acutely isolated hippocampal LD-lo and LD-hi microglia from aged mice and measured cytokine concentrations in the supernatant using multiplex array 8h after stimulation with LPS or saline as a control. We found that under baseline conditions (saline treatment), LD-hi microglia released increased levels of multiple cytokines, including CCL3, CXCL10, and IL-6, compared to LD-lo microglia. In addition, LD-hi microglia showed a strongly exaggerated release of multiple cytokines, such as IL-10, CCL3, CCL4, IL-6, CCL5, TNF-α IL-1β IL-1α CXCL1 and CXCL10 upon LPS treatment compared to LD-lo microglia (Fig. 5i,j). These findings suggest that LAM are in a primed activation state that becomes hyperactivated upon stimulation with LPS.

### CRISPR-Cas9 screen identifies genes linked to neurodegeneration as genetic regulators of lipid droplet formation

To investigate genetic regulators of microglial lipid droplet formation, we performed pooled CRISPR-Cas9 screens. Informed by our RNAseq data of LAM (Fig. 2c,e), we chose to screen with an sgRNA library targeting ∼ 2000 genes involved in the lysosomal pathway and protein degradation as well as in cellular stress, with 10 distinct sgRNAs targeting each gene along with ∼1000 negative control sgRNAs^53^. To probe the role of these genes in lipid droplet formation, we used the microglial BV2 cell line to generate a pooled population of targeted BV2 cells for every gene represented in the sgRNA library. LPS was used to induce lipid droplets in these cells.

To identify sgRNAs that inhibited or promoted the formation of lipid droplets, we developed a photoirradiation selection strategy in which BV2 cells are separated based on their capacity to form lipid droplets. By adding iodine atoms to the lipid droplet marker BODIPY(iodo-BODIPY, iBP; the synthesis followed the procedure described previously by Nagano and coworkers^54^), we transformed this molecule into a photosensitizer that induces cell death in iBP^+^ cells after photoexcitation^54, 55^ (Fig. 6a). To prove the efficacy of the photoablation approach we used Calcein live cell imaging and found that irradiation of iBP^+^ BV2 cells selectively killed lipid-droplet rich cells (Fig. 6b,c).

**Fig. 6.**
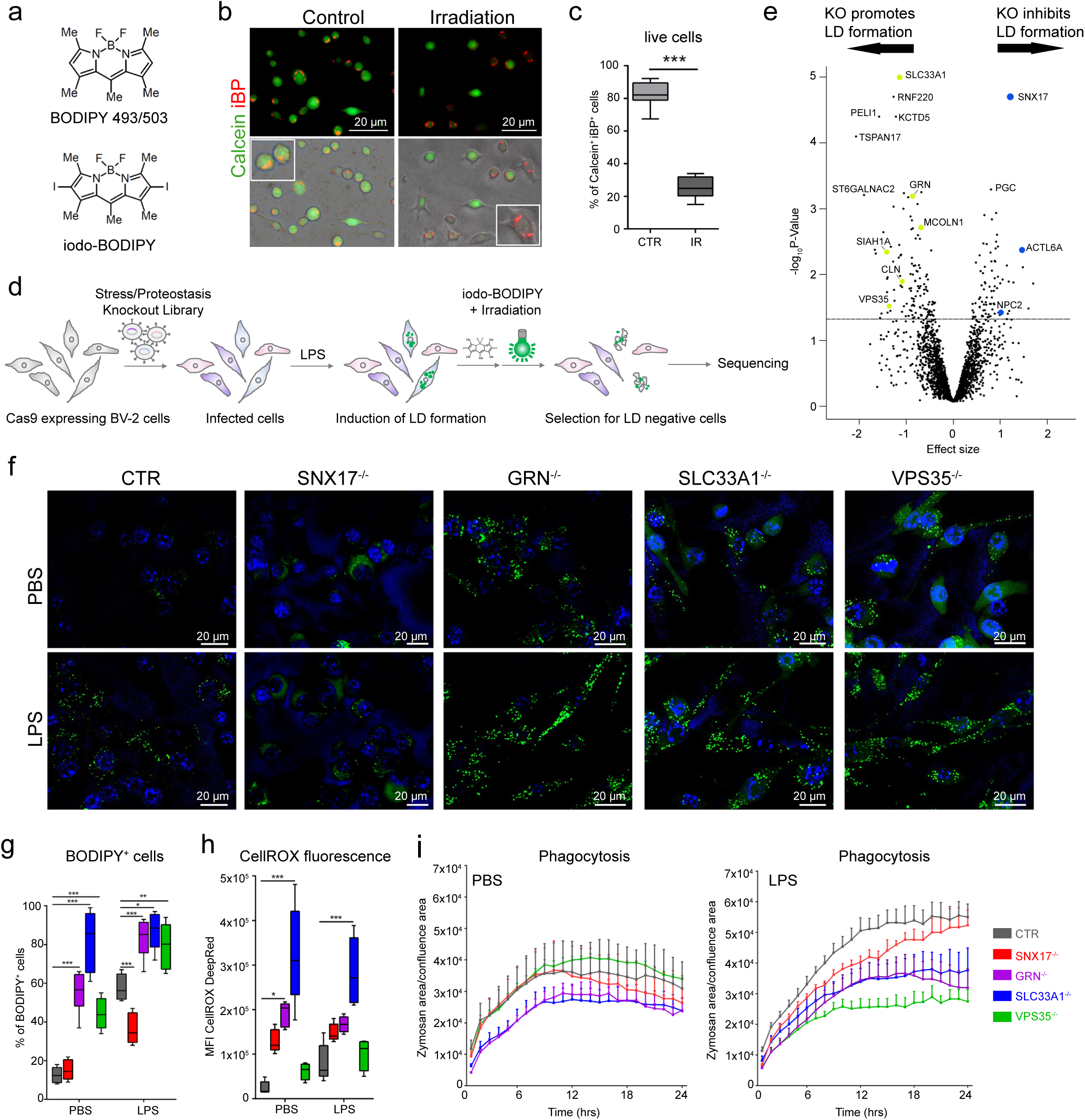
CRISPR-Cas9 screen identifies genetic regulators of lipid droplet formation. **a**, Structure of BODIPY 493/503 and of iodo-BODIPY. **b**,**c**, Calcein^+^ signal in BV2 cells that were treated with 5 μg/ml LPS for 18 hours, stained with iBP, and exposed to photoirradiation for 3 hours. Representative micrographs (**b**) and quantification (**c**) of Calcein^+^ (live cells) and iBP^+^ (lipid-droplet containing cells) in control (non-irradiated) and irradiated BV2 cells. **d**, Experimental schematic for pooled CRISPR-Cas9 screen to identify regulators of lipid droplet formation in LPS-treated BV2 cells. **e**, Volcano plot showing hits for genetic regulators of lipid droplet formation from the CRISPR-Cas9 knockout screen. Dashed line represents P-value < 0.05 cutoff. Positive effect size represents genes targeted by sgRNAs that were enriched in lipid droplet-negative cells; negative effect size represents genes targeted by sgRNAs that were under-represented in lipid droplet-negative cells. Genes previously associated with neurodegeneration are highlighted in blue and yellow. **f-i**, Single CRISPR-Cas9 knockout BV2 cell lines of selected screen hits (*SNX17*^-/-^, *GRN*^-/-^, *SLC33A1*^-/-^, *VPS35*^-/-^). Cas9-expressing BV2 cells were used as control (CTR). Representative micrographs of BODIPY^+^ staining in PBS or LPS treated cells (**f**) and quantification of the percentage of BODIPY^+^ cells (**g**). Quantification of CellROX MFI in PBS or LPS treated cells (**h**). Time lapse imaging and quantification of Zymosan uptake in cells treated with PBS or LPS (**i**). Experiments on BV2 cells were performed three times in technical triplicates. Statistical tests: unpaired Student’s t-test (c), two-way ANOVA (g,h,i) followed by Tukey’s post hoc test. Error bars represent mean ± SD (c,g,h) and mean ± SEM (i). Horizontal lines in the box plots indicate medians, box limits indicate first and third quantiles, and vertical whisker lines indicate minimum and maximum values. *P<0.05, **P< 0.01, ***P< 0.001. LD, lipid droplet; iBP, iodo-BODIPY; MFI, mean fluorescent intensity.

After three rounds of selection against lipid droplet-rich cells we sequenced the sgRNA composition of the selected lipid droplet-negative BV2 cells. We found 112 genes that were significant positive or negative regulators of lipid droplet formation (P<0.05; FDR<50%; Fig. 6e). Unexpectedly, the top hits included various genes that have been previously linked to neurodegeneration including *SLC33A1*, *SNX17*, *GRN* and *VPS35*^56–59^, hinting at a possible relationship between lipid storage in microglia and neurodegeneration.

We generated individual BV2 cell lines with CRISPR deletions of selected hits that were detected as negative (*SLC33A1*, *GRN, VSP35*) or positive (*SNX17*) regulators of lipid droplet formation in the screen. Indeed, BV2 cells with sgRNAs targeting *GRN*, *SLC33A1* and *VPS35* had significantly more lipid droplets than BV2 cells with control sgRNAs, and sgRNAs targeting *SNX17* inhibited lipid droplet formation upon LPS treatment (Fig. 6f,g). We used flow cytometry to measure ROS levels and found significantly increased ROS generation in BV2 cells with sgRNAs targeting *GRN* and *SLC33A1* cells under baseline conditions and in *SLC33A1* sgRNA expressing cells upon LPS treatment (Fig. 6h). Next, we assessed phagocytosis by analyzing pHrodo red Zymosan uptake and found that cells with sgRNAs targeting *GRN* and *SLC33A1* had significant deficits in Zymosan uptake compared to control cells. In *VPS35* sgRNA expressing cells, phagocytosis was specifically compromised upon LPS treatment but not under baseline conditions (Fig. 6i).

### GRN^-/-^ mice contain high numbers of lipid droplet-accumulating microglia which have functional impairments and a transcriptional signature similar to LAM

To confirm the findings from our screen *in vivo*, we analyzed lipid droplet numbers in microglia in *GRN*^-/-^ mice. *GRN* mutations are linked to the development of frontotemporal dementia (FTD)^58^, and *GRN*^-/-^ mice are used as a model for FTD and are characterized by microglial changes, neuroinflammation, and cognitive deficits^60^. Given the results of our CRISPR-Cas9 KO screen, we hypothesized that the *GRN* KO would promote lipid droplet formation. We used middle-aged (9-10 months) mice, because at this age GRN^-/-^ mice already show behavioral changes and brain impairments^60^, while wild type mice present only minor signs of neuroinflammation and contain low numbers of LAM. We found that microglia in the hippocampus of *GRN*^-/-^ mice contained high numbers of lipid droplets, resulting in a twofold higher percentage of BODIPY^+^ microglia and twice as many lipid droplets per cell in GRN^-/-^ mice compared with wild type littermates (Fig. 7a-c). Moreover, we frequently detected BODIPY^+^ Iba1^+^ microglia in the thalamus and occasionally also in the cortex and corpus callosum in GRN^-/-^ mice (Supplementary Figure 5).

**Fig. 7.**
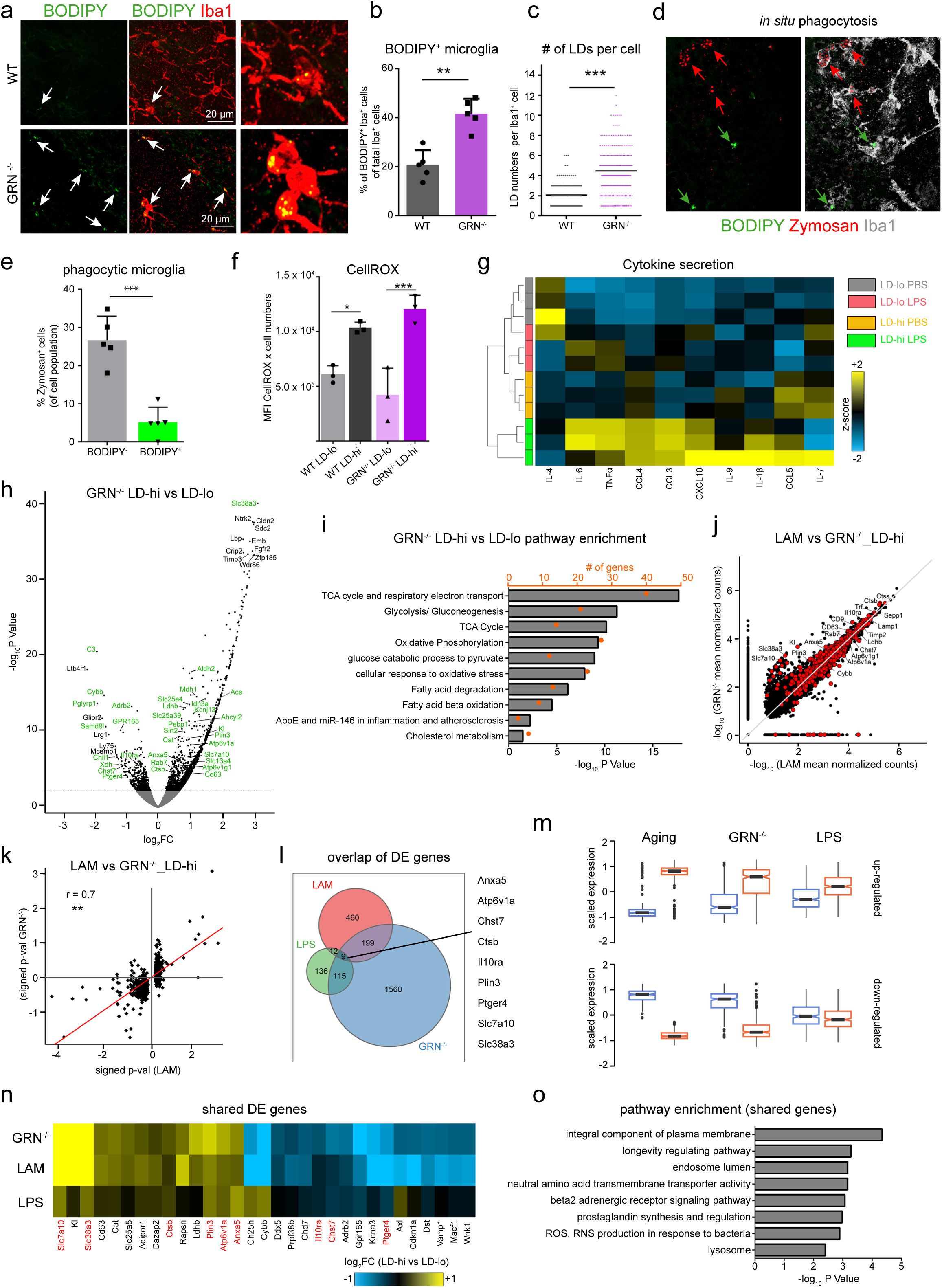
GRN^-/-^ mice possess high numbers of lipid droplet-rich microglia which functionally and partially transcriptionally resemble LAM. **a-c**, BODIPY^+^ and Iba1^+^ expression in the hippocampus of 9-month old male WT mice and age- and sex-matched *GRN*^-/-^ mice. Representative confocal images (**a**), and quantification of BODIPY^+^/Iba1^+^ microglia (**b**) and of lipid droplet numbers per Iba1^+^ microglia (**c**). n = 5 mice per group. **d**,**e**, 250 μm organotypic brain slices from 9-month old GRN^-/-^ mice were incubated for 4 hours with pHrodo red Zymosan particles. Representative confocal images of the hippocampus (**d**) and quantification of Zymosan uptake in BODIPY^-^ and BODIPY^+^ Iba1^+^ cells (**e**). n = 3 mice per group. **f**, Quantification of CellROX fluorescence in primary LD-lo and LD-hi microglia from 9-month old GRN^-/-^ mice and age-matched wild type controls. **g**, Acutely isolated LD-lo and LD-hi primary microglia from 10-month old GRN^-/-^ mice were treated with LPS (100 ng/ml) or PBS for 8 hours. Heatmap shows changes in cytokine secretion under baseline conditions (PBS) and after LPS treatment. **h**-**k**, RNA-Sequencing of BODIPY^lo^ (=LD-lo) and BODIPY^hi^ (=LD-hi) CD11b^+^CD45^lo^ microglia from the hippocampus of 10-month old GRN^-/-^ mice. n = 3 samples per group. Each sample is a pool of microglia from 2 mouse brains. **h**, Volcano plot showing differentially expressed genes in LD-hi versus LD-lo microglia. Dashed line represents q-value < 0.05 cutoff. LAM genes are highlighted in green. **i**, EnrichR pathway analysis of genes differentially regulated between LD-hi and LD-lo microglia. **j**, Scatterplot showing gene expression intensities (mean normalized counts) of LD-hi microglia in GRN^-/-^ mice compared to LAM in aging. Genes differentially expressed in LAM are highlighted in red. **k**, Scatterplot showing expression changes of LAM DE genes in LD-hi microglia from GRN^-/-^ mice and in LAM in aging. **l**, Overlap of genes differentially expressed between LD-hi and LD-lo microglia in aging (LAM), in GRN^-/-^ mice, and in LPS-treated young mice, and list of genes that are shared between all groups. **m**, Box plots showing the scaled expression of up- and downregulated DE LAM genes (692 genes) in aging, GRN^-/-^ mice, and LPS-treated young mice. **n**,**o**, Heatmap showing expression changes (**n**) and EnrichR pathway analysis (**o**) of the shared genes (red) and the 20 most significant genes shared between LAM and LD-hi microglia in GRN^-/-^ mice (ranked by P-value). Experiments on primary cells (f,g) were performed two times in technical triplicates. Statistical tests: unpaired Student’s t-test (b,c,e) and one-way ANOVA (f) followed by Tukey’s post hoc test. Error bars represent mean ± SD. *P<0.05, **P< 0.01, ***P< 0.001. DE genes, differentially expressed genes; LD, lipid droplet; MFI, mean fluorescent intensity.

To determine whether lipid droplet containing microglia from GRN^-/-^ mice have functional defects similar to LAM, we measured phagocytic activity, ROS production, and cytokine secretion. Indeed, BODIPY^+^ microglia in GRN^-/-^ mice showed decreased uptake of pHrhodo Zymosan in an *in situ* slice culture phagocytosis assay (Fig. 7d,e), and LD-hi microglia isolated from GRN^-/-^ mice showed increased levels of CellROX fluorescence in the ROS assay, as well as elevated secretion of pro-inflammatory cytokines after stimulation with LPS, compared to LD-lo GRN^-/-^ microglia (Fig. 7f,g).

Next, we asked whether LD-hi microglia from GRN^-/-^ mice are transcriptionally similar to LAM from aged mice. We performed RNA-sequencing on LD-lo and LD-hi microglia isolated from 10-month-old GRN^-/-^ mice and found 1884 differentially expressed genes (Fig. 7h). The most significant pathways associated with these genes included metabolic processes and cellular response to stress (Fig. 7i), pathways that are also enriched in LAM (Fig, 2e, Supplementary Fig. 2b). To compare the gene expression patterns of LD-hi GRN^-/-^ microglia and of LAM in aging, we plotted expression intensities (Fig. 7j) and expression changes (Fig.7k) of genes differentially expressed in LAM and found a strong correlation between both datasets.

Finally, we sought to explore the transcriptional commonalities of lipid droplet-rich microglia in aging, in GRN^-/-^ mice, and in LPS-treated young mice. We found that although only nine common genes were differentially expressed in all three data sets (one of them was the lipid droplet specific gene *PLIN3*), the directionality of expression changes of LAM genes was remarkably similar for all groups and almost identical for LAM and LD-hi GRN^-/-^ microglia (Fig. 7l,m; Supplementary Figure 6). To identify enriched pathways for genes shared between lipid droplet-rich microglia, we used the nine genes shared between all data sets and additionally included the top 20 common genes between LAM and LD-hi GRN^-/-^ microglia, because of their high transcriptional similarity. Interestingly, significant pathways included *endosome lumen*, *ROS production*, and *lysosome*, pathways linked to the dysfunctional phenotype observed in LAM and in LD-hi GRN^-/-^ microglia (Fig. 7n,o).

Together, these findings validate the *in vivo* relevance of our *in vitro* CRISPR screen and confirm that microglia from GRN^-/-^ mice contain lipid droplets. Remarkably, LD-hi microglia from GRN^-/-^ mice and LAM in aging have similar functional and transcriptional phenotypes, suggesting that lipid droplet containing GRN^-/-^ microglia share the LAM state.

## Discussion

In this study, we identify a novel, lipid droplet accumulating microglia (LAM) state in the aging brain and show that LAM are characterized by a unique transcriptional signature, severe functional deficits and a pro-inflammatory phenotype.

### What is the role of LAM in the aging brain?

During aging, microglia undergo profound transcriptional and functional changes. They are considered to be in a “primed” state and show an increased baseline production of pro-inflammatory cytokines such as TNF-α, IL-1β and IL-6, and become hyper-activated upon immune challenge^48^. Further, they are compromised in their phagocytosis activity and produce elevated levels of reactive oxygen species (ROS)^5, 7, 8, 61^. These microglia changes are suggested to play a vital role in age-related neuroinflammation and in structural and functional impairments in the aging brain.

Microglia in the LAM state, which account for more than 50% of all microglia in the aged hippocampus, are impaired in phagocytosis, generate elevated levels of ROS, produce high amounts of pro-inflammatory cytokines under resting conditions and react with excessive cytokine release upon immune challenge. These findings demonstrate that LAM, but not microglia without lipid droplets, show typical age-related functional impairments and a primed phenotype, suggesting that LAM is a detrimental microglia state in the aging brain.

The idea that LAM have an unfavorable role in brain aging is in line with previous findings which have shown that lipid droplet containing immune cells in the periphery are detrimental for their environment. For example, it has been shown that lipid droplet accumulating foamy macrophages in atherosclerosis are deleterious for disease pathology, and pharmacological inhibition of lipid droplet formation decreased atherosclerotic areas and had anti-inflammatory effects^26^. Similarly, lipid droplet rich eosinophils in experimental models of allergy are associated with an exacerbated inflammatory response and strategies that inhibit lipid droplet formation in eosinophils may be therapeutically useful in controlling allergic inflammation^23^. However, lipid droplets have also been correlated with beneficial functions, such as improved host defense and antigen cross presentation in myeloid cells^20^. In the brain, a protective role has been reported for lipid droplets in glia in a drosophila model for neurodegeneration^30, 31^. Hence, whether lipid droplet containing cells are beneficial or detrimental for their milieu might be dependent on the cell type and the environmental context. Here, we show that LAM represent a dysfunctional, “primed” state of microglia, and suggest that LAM are harmful for the aging brain. Ultimately, pharmacological and genetic ablation experiments will be required to prove the role of LAM in brain aging and disease.

### What causes lipid droplet formation in LAM?

Lipid droplets can form due to various environmental and cellular conditions, including elevated concentrations of extracellular lipids, inflammatory events, increased ROS levels and intracellular metabolic changes^62–64^. In peripheral immune cells, lipid droplets form often as a response to inflammation and stress^20, 22^. Interestingly, we found that LPS treatment, which provokes an acute inflammatory response in the brain, led to lipid droplet formation in hippocampal microglia in young mice *in vivo* and in the microglia cell line BV2 *in vitro*. Furthermore, we found lipid droplet containing microglia in GRN^-/-^ mice, a model for frontotemporal dementia that is characterized by severe chronic neuroinflammation. Here, lipid droplet containing microglia were not restricted to the hippocampus but also frequently found in the thalamus. This is of particular interest because it has been shown that in GRN^-/-^ mice, neuroinflammation is most pronounced in the thalamus^83^. These findings are corroborated by previous reports about lipid droplets in microglia, which concordantly observed lipid droplet formation under inflammatory or stress conditions. Specifically, lipid droplets in microglia have been detected in rat brains after stroke^34^, in adult mice after induction of ROS by rotenone^30^, in hippocampal slice culture from P6-8 mice upon treatment with LPS, as well as in LPS-treated microglia from mixed neural cultures and in LPS-treated cells of the microglia-like cell line N9^35,65^.

Thus we suggest that inflammation plays a key role in the buildup of lipid droplets in microglia, and it is tempting to speculate that age-related neuroinflammation provokes the formation of LAM. Similar to foamy macrophages in atherosclerosis and to lipid droplet accumulating eosinophils in allergy, LAM might occur in an inflamed environment, and subsequently trigger an exacerbated inflammatory response. Future investigations will show whether strategies that decrease neuroinflammation in aging or in neurodegenerative diseases are able to inhibit or reverse lipid droplet accumulation in microglia.

Besides inflammation, metabolic changes towards increased fatty acid production have been reported to cause lipid droplet formation in immune cells^20, 66, 67^ and also in cancer cells^68^. Interestingly, genes associated with the *fatty acid beta oxidation* pathway are significantly upregulated in LAM. For example, expression of *ACLY*, a key enzyme in the shift from the TCA cycle to lipid formation that has been shown to cause lipid droplet accumulation^69^, is significantly higher in LAM than in microglia without lipid droplets. Similarly, RNA-Seq analysis of lipid droplet-rich microglia in LPS-treated young mice and in GRN^-/-^ mice revealed significant enrichment of pathways related to metabolism, including *TCA cycle* and *fatty acid beta oxidation*. Moreover, LAM show a significantly higher NAD^+^/NADH ratio compared to microglia without lipid droplets, implying metabolic alterations in these cells (Supplementary Fig. 6). These findings suggest increased lipid synthesis in LAM, yet it remains to be shown to which extent metabolic changes are involved in microglial lipid droplet formation in aging.

Cholesterol accumulation has recently been observed in phagocytes in an EAE demyelination model in aged mice as a result of excessive uptake of myelin debris^70^. Because demyelination is a characteristic of brain aging, we analyzed the cholesterol content of lipid droplets in LAM. We did not find ultrastructural evidence for cholesterol crystals, and lipidomics analysis revealed that these lipid droplets contained mainly glycerolipids and only low amounts of cholesteryl ester. Moreover, the lipid composition of lipid droplets from young microglia and from old microglia was almost identical (Supplementary Fig. 2). Thus, we suggest that demyelination does not contribute to lipid droplet formation in LAM.

In a drosophila model for neurodegeneration^30, 31^, lipid droplets in glia form due to an APOE dependent transfer of lipids from neurons to glia. Given that transcription of APOE and other lipid transporters was either down- or not significantly regulated in LAM (Supplementary Table 1), it is unlikely that this mechanism leads to lipid droplet formation in aged microglia.

Collectively, these findings support the idea that lipid droplets in LAM form due to the inflammatory environment in the aging brain, rather than due to extracellular uptake of lipids such as myelin or neuron-derived lipids. Whether a pro-inflammatory environment induces metabolic changes in aged microglia which subsequently lead to lipid droplet formation remains to be clarified.

### Role of lipid droplets in the LAM functional phenotype

Increased ROS generation is a main characteristic of LAM and is also observed in peripheral lipid droplet containing immune cells. Reports about whether ROS is a cause or consequence of lipid droplet formation are contradictory^71–73^. Interestingly, our *in vitro* results demonstrated that pharmacological inhibition of lipid droplet formation with Triacsin C prevented ROS generation in BV2 cells, supporting the idea that triglycerides and lipid droplets have a causal role in the LPS-induced generation of ROS. It remains to be deciphered how lipid droplets contribute to increased ROS formation in LAM in the aging brain. Similar to the idea that neuroinflammation is both a cause and consequence of lipid droplet formation in LAM, it is possible that elevated ROS initially trigger lipid droplet formation, and subsequently lipid droplets induce ROS formation and exacerbate intracellular ROS load.

LAM showed severe phagocytosis deficits compared to microglia without lipid droplets in the aging brain. This finding is in line with a previous report which observed that at the sites of atherosclerotic lesions, lipid droplet rich foamy macrophages show decreased phagocytosis activity compared to macrophages without lipid droplets^74^. Again, defective phagocytosis could be a cause or consequence of lipid droplet accumulation. Our *in vitro* findings showed that pharmacological inhibition of lipid droplet formation could significantly increase phagocytosis in BV2 cells, suggesting a detrimental role of lipid droplets for phagocytosis. The exact mechanism of how lipid droplets might interfere with phagocytosis remains to be shown. In this context, a study in macrophages has shown that engulfment of cellular debris relies on the availability of free fatty acids, which are released upon degradation of lipid droplets, thus linking effective lysis of lipid droplets with successful phagocytosis^75^. Notably, RNA-Seq analysis revealed that *ADRB1* and *ADRB2*, two key enzymes in the process of lipid degradation, are significantly downregulated in LAM. This suggests impaired lysis of lipid droplets in LAM, and subsequent lack of free fatty acids might be an underlying reason for impaired phagocytosis in these cells.

In addition to enzymatic lysis, lipid droplets can be degraded by lysosomes, a process termed “lipophagy”^76^. Intriguingly, we observed that LAM contain high numbers of lysosomes, and moreover, these lysosomes accumulate in close contact to lipid droplets. It is possible that in LAM, lysosomes are used for lipophagy of lipid droplets rather than for degrading phagocytosed material, resulting in impaired phagocytosis. Lastly, there is emerging evidence that lysosomes become dysfunctional in aged microglia^5^. Hence, there is the possibility that in LAM, lipid droplets accumulate due to defective degradation processes, and lipid droplet accumulation and impaired phagocytosis could be two co-existing yet independent processes that are the consequence of defective lysosomes.

### LAM in neurodegeneration

Recently, it has been shown that several subsets of microglia with unique molecular and functional phenotypes exist in the healthy, aged, and degenerating brain^10–12^. For example, Keren-Shaul and co-workers^12^ (2017) identified a protective, phagocytic microglia subtype (“DAM”) with the potential to restrict neurodegeneration, and Krasemann and colleagues^11^ (2017) identified a dysfunctional, neurodegenerative microglia phenotype (“MGnD”). The LAM transcriptional signature showed almost no overlap with DAM and MGnD, and surprisingly, typical aging genes^40^ such as *AXL*, *CLEC7A* and *CYBB*, are regulated in a reciprocal direction in LAM. Furthermore, *TREM2* and *APOE*, two key genes involved in the progression of neurodegeneration that are upregulated in DAM and MGnD, are not regulated in LAM. Likewise, LAM showed only low overlap with the transcriptome of recently identified “lipid-associated macrophages” from mouse and human adipose tissue, a cell population that controls metabolic homeostasis in a *TREM2* dependent manner and is transcriptionally similar to DAM^84^. LAM and DAM also show different functional phenotypes; while DAM are actively phagocytic populations, LAM are severely impaired in this function. Thus, we suggest that LAM represent a state distinct from DAM and MGnD in the aging brain. The specific dynamics of the LAM state in aging and disease remain to be deciphered, and it remains to be shown whether microglia subtypes such as DAM and MGnD transform into the LAM cellular state at some point during disease progression.

By using pooled CRISPR-Cas9 targeted screening, we discovered genes for which variants cause autosomal dominant forms of neurodegeneration (*SLC33A1*, *SNX17*, *GRN, VPS35, CLN, NPC2*) as genetic modulators of lipid droplet formation in microglia^56–59, 77^. Indeed, sgRNAs targeting these genes were able to increase (*SLC33A1*, *GRN*, *VPS35*) and decrease (*SNX17*) lipid droplet load in microglial BV2 cells. In addition, sgRNAs targeting *GRN* and *SLC33A1* induced significant defects in phagocytosis and increased ROS production in BV2 cells, hence recapitulating the functional impairments of LAM.

Excitingly, knockout of *GRN*, which causes frontotemporal dementia in people with loss of function mutations in this gene^58^, resulted in severe accumulation of lipid droplets in microglia *in vivo*. These lipid droplet rich microglia in GRN^-/-^ mice show a similar transcriptome signature and the same functional impairments (reduced phagocytosis, increased ROS formation, excessive secretion of proinflammatory cytokines) as LAM in aging, suggesting that they share the LAM state. The finding of lipid droplet containing microglia in GRN^-/-^ mice is corroborated by a recent study which showed that cultured progranulin-deficient macrophages exhibited increased foam cell formation, i.e. lipid droplet accumulation, compared to wild type macrophages^78^. In addition, a recent study reported that loss of *GRN* leads to intracellular and intra-lysosomal accumulation of long polyunsaturated triacylglyerides^79^ in the brains of humans and mice. Since triacylglycerides (TAG) are a major component of lipid droplets in LAM, it is possible that lipid droplet accumulation in GRN deficient microglia contributes to elevated TAG levels in brains lacking GRN. It has to be shown to what extent lipid droplet containing microglia contribute to disease pathology in frontotemporal dementia.

Together, we show that LAM demonstrate a novel state of microglia with a unique transcriptional signature and functional impairments in the aging brain, and we identified lipid droplet containing microglia in a GRN^-/-^ model for chronic neuroinflammation, and in an LPS-induced acutely inflamed brain milieu. Future studies will show whether LAM are a common characteristic for neuroinflammation, and whether they have a role in neurodegenerative diseases.

In the future, targeting LAM might represent an attractive and druggable approach to decrease neuroinflammation and to restore brain homeostasis in aging and neurodegeneration, with the goal to improve cognitive functions.

## Author Contributions

J.M. and T.W.-C. conceptualized and designed the study, analyzed and interpreted data, and wrote the manuscript. J.M. and S.E.L designed the figures. J.M., T.I., M.Z., S.E.L., and J.V.P. acquired the data. J.M. performed electron microscopy. J.M. and M.Z. performed histology and organotypic slice culture experiments. J,M, T.I. and M.Z. performed cell culture experiments. J.M., T.I. and S.E.L performed RNA Seq experiments. J.V.P., M.Z. and J.M. performed stereotactic procedures. V.M. conducted in vivo LPS injections and provided GRN^-/-^ mouse brain sections. J.M., M.S.H. and D.W.M. generated and analyzed CRISPR-Cas9 screen data. J.M. and B.L. analyzed RNA-seq data. J.T.,T.F. and O.H. performed mass-spectrometric experiments. J.M. and H.W. performed CARS imaging. J.K. and C.B. designed and produced methylated BODIPY derivatives. All authors reviewed the manuscript.

## Acknowledgements

We thank the members of the Wyss-Coray laboratory for feedback and support throughout the study, Prof. Walter Stoiber (University of Salzburg, Austria) and the Division of Optical and Electron Microscopy, University of Salzburg, Austria, for excellent assistance with EM images, and Divya Channappa from the Department of Neurology and Edward Plowey from the Department of Pathology, Stanford University, for providing human post mortem brain samples. CARS imaging was performed at the Microscopy Core Facility of the Institute of Molecular Biosciences, University of Graz, Austria. We thank Prof. Annika Enejder, Heilshorn Biomaterials Group, Stanford University for discussion and for reviewing the manuscript. This work was supported by the FWF Hertha-Firnberg Postdoctoral program n° T736-B24 (J.M.), the PMU-FFF E-16/23/117-FEA (T.K.F), the Stanford Neuroscience Institute Brain Rejuvenation Project Award and NIH Director’s New Innovator Award (1DP2HD084069-01) to M.C.B., the Department of Veterans Affairs (T.W.-C.), the National Institute on Aging (DP1-AG053015 to T.W.-C.), the NOMIS Foundation (T.W.-C.), the Glenn Foundation for Medical Research (T.W.-C.), the Cure Alzheimer’s Fund (T.W.-C.) and the Nan Fung Life Sciences Aging Research Fund (T.W.-C.).

---

**Supplementary Fig. 1 Lipid droplet accumulating microglia are abundant in the hippocampus but rare in other brain regions of aged mice.** *Related to Figure 1* a-d, Representative confocal images of the cortex (**a**), thalamus (**b**), corpus callosum (**c**) and hippocampal dentate gyrus (**d**) from 20-month old male mice stained for BODIPY^+^ (lipid droplets) and Iba1^+^ (microglia). Scale bar: 20 μm. Arrows point towards BODIPY^+^ lipid droplets. e, Quantification of BODIPY^+^/Iba1^+^ cells. n = 4 mice per group. One-way ANOVA followed by Tukey’s post hoc test. Error bars represent mean ± SD. ***P< 0.001.

**Supplementary Fig. 2 LAM have a unique transcriptional signature that minimally overlaps with published gene expression profiles of microglia in aging and neurodegeneration.** *Related to Figure 2* a,**b**, IPA pathway analysis of genes that are significantly upregulated (a) or downregulated (b) in LD-hi microglia in aging. Analysis based on top 100 down- and up-regulated genes. **c**-**g**, Expression plots comparing RNA-Seq data of LAM (see Fig. 2) with published RNA-Seq data of microglia in aging (**c**), AD (**d**), ALS (**e**), disease-associated microglia (DAM) (**f**) and neurodegenerative microglia (MGnD) (**g**). Data are expressed as signed fdr, i.e the product of log2 FC and log10 fdr. **h**, Paired dot plot showing FPKM values of LD-lo and LD-hig microglia for ApoE (P= 0.423). Dotted lines connect LD-lo and LD-hi microglia sorted from the same samples. **i**, Heatmap showing expression changes of LAM genes (genes differentially expressed in LD-hi microglia in aging) in LD-hi microglia from GRN^-/-^ mice, from LPS treated mice, and in microglia clusters revealed by Li et al. (2019) and Hammond et al. (2019)^14, 15^. LD, lipid droplet.

**Supplementary Fig. 3 LPS treatment induces lipid droplet formation in microglia and in BV2 cells.** *Related to Figure 3* **a**,**b**, 3-month-old male mice were given intraperitoneal (i.p.) injections of LPS (1 mg/kg BW) for four days. Representative confocal images of BODIPY^+^ and Tmem119^+^ in the hippocampus (**a**) and of BODIPY and Iba1 staining in the cortex, corpus callosum, and thalamus (**b**). **c**-**e**, Lipidome profiling of lipid droplets from LPS-treated BV2 cells, primary microglia, and liver tissue. **c**, Pie charts showing that the lipid composition of lipid droplets from young and aged microglia is highly similar, but differs between young and aged liver tissue. **d**,**e**, Distribution of MAG chain lengths (**d**) and TAG saturation levels (**e**) of lipid droplets isolated from LPS-treated BV2 cells and from microglia and liver tissue from aged mice. young= 5-month-old male mice, old= 20-month-old male mice; n = 4 mice per group.

**Supplementary Fig. 4 Aged plasma induces lipid droplet formation in BV2 cells.** *Related to Figure 4* **a**, Representative micrographs of BODIPY^+^ staining and of phagocytosis of of pHrodo red Zymosan in BV2 cells treated with 5% plasma from young (3-months) and aged (18-months) mice for 12 hours. **b**, Quantification of BODIPY^+^ staining in BV2 cells treated with young and aged plasma. **c**,**d,** Quantification of Zymosan uptake in BV2 cells treated with young and aged plasma (**c**), and in aged plasma treated BODIPY-low and BODIPY-high cells (**d**). Statistical tests: unpaired Student’s t-test. Error bars represent mean ± SD. *P< 0.05, ***P< 0.001.

**Supplementary Fig. 5 Lipid droplet containing microglia in the cortex, corpus callosum, and thalamus of GRN^-/-^ mice.** *Related to Figure 7* **a**-**c**, Representative confocal images of BODIPY^+^ (lipid droplets) and Iba1^+^ (microglia) in the cortex (**a**), corpus callosum (**b**), (**c**) and thalamus from 9-month-old male GRN^-/-^ mice. BODIPY^+^/Iba1^+^ cells were frequently found in the thalamus and were detected to a lesser extent in cortex and corpus callosum.

**Supplementary Fig. 6 Expression changes of LAM genes in lipid droplet-rich microglia from normal aging, GRN^-/-^ and LPS-treated mice.** *Related to Figures 3 and 7* a, Heatmap showing expression changes of LAM genes (genes differentially expressed in LD-hi microglia in aging; 692 genes) in LD-hi microglia from GRN^-/-^ mice and from LPS treated mice.

**Supplementary Fig. 7 LAM show signs of metabolic alterations.** *Related to Figure 2* **a**, Paired dot plot showing FPKM values of LD-lo and LD-hi microglia for ACLY (data obtained from RNA-Seq analysis, see Figure 2). Dotted lines connect LD-lo and LD-hi microglia sorted from the same samples. **b**, NAD colorimetric assay showing the NAD^+^/NADH ratio of primary hippocampal microglia from 3-month old mice (young) and of LD-lo and LD-hi primary microglia from 20-month old male mice. Experiments were performed two times in technical triplicates. n=3 mice per group per experiment. Statistical tests: one-way ANOVA (b) followed by Tukey’s post hoc test. Horizontal lines in the box plots indicate medians, box limits indicate first and third quantiles, and vertical whisker lines indicate minimum and maximum values. *P< 0.05, ***P< 0.001.

**Supplementary Table 1 List of genes previously related to lipid droplet biology.** *Related to Figures 2,3,6,7* RNA-Seq and CRISPR-Cas9 genetic screen results for lipid droplet-related genes. Genes that were significantly regulated in the RNA-Seq analysis in normal aging (Fig. 2), in LPS-treated young mice (Fig. 3), in GRN^-/-^ mice (Fig. 7), or in the CRISPR screen are highlighted in red.

**Supplementary Table 2 RNA-Seq data, CRISPR-Cas9 genetic screen results, IPA pathway analysis, and gene lists from published microglia data sets.** *Related to Figures 2,3,6,7*

## METHOD DETAILS

### Animals

Aged C57BL/6J male wild type mice (18-20 months old) were obtained from the National Institute on Aging (NIA), and young C57BL/6J males (2-4 months of age) were purchased from Jackson Laboratory. Grn^-/-^ deficient mice (B6.129S4(FVB)-Grntm1.1Far/Mmja) and wild type littermates were bred and aged in-house but originally acquired from Jackson. Mice were housed under a 12-hour light-dark cycle in pathogenic-free conditions, in accordance with the Guide for Care and Use of Laboratory Animals of the National Institutes of Health. All animal procedures were approved by the V.A. Palo Alto Committee on Animal Research and the institutional administrative panel of laboratory animal care at Stanford University. Male mice were used for all experiments.

### LPS injections

3-month-old male wild type mice were injected with LPS (Lipopolysaccharide from *E. coli*, Sigma), i.p. 1 mg LPS/kg body weight once daily for four consecutive days. Control mice were injected with body weight corresponding volumes of PBS. 24 hours after the last LPS injection mice were euthanized, brains were extracted and brain tissue was processed for immunohistochemistry staining (see *Perfusion and tissue processing* and *Immunohistochemistry and BODIPY staining*) or for microglia isolation (see *Microglia Isolation*).

### Electron Microscopy

3- month and 20-month-old male C57BL/6 wild type mice (n=3 per group) were anesthetized with 3.8% chloral hydrate (wt/vol) and transcardially perfused with 0.9% saline, followed by 2% paraformaldehyde/ 2.5% glutaraldehyde in 0.1M phosphate buffer (PB; pH7.4). Brains were removed and postfixed in the same fixative overnight (4°C) and then stored in 0.1 M PB. Sagittal 200 µm sections were cut on a Leica VT1000S vibratome (Leica). The sections were incubated in 2% osmium (Electron Microscopy Sciences) for 2 h, rinsed in 0.1M PB, dehydrated in a graded series of ethanol and embedded in Araldite (Durcupan, Electron Microscopy Sciences). Semithin (1.5 µm) and ultrathin (70-80 nm) sections of the dentate gyrus were cut using a Reichert Om-U 3 ultramicrotome (Leica). Ultrathin sections were mounted on Formvar coated 75-mesh copper grids, contrasted with aqueous solutions of uranyl acetate (0.5%) and lead citrate (3%), and analyzed at 80 kV in a EM 910 transmission electron microscope (Zeiss) equipped with a Troendle sharp:eye 2k CCD camera.

To evaluate the ultrastructure of microglia within the dentate gyrus, 10 ultrathin sections were analyzed per mouse. Microglia were identified by a combination of ultrastructural characteristics, including a highly electron-dense cytoplasm and nucleus, an often star-shaped cell morphology, an irregularly shaped nucleus with coarsely clumped chromatin, and a cytoplasm rich in free ribosomes and vesicles^1^. Cytoplasm and nucleus area were analyzed using ImageJ software 1.45s.

### Perfusion and tissue processing

Mice were anesthetized using Avertin (Tribromoethanol) and transcardially perfused with 0.9% NaCl solution. Brains were extracted, fixed in 4% paraformaldehyde for 48 hours, cryoprotected in 30% sucrose and then sectioned sagitally or coronally (40 μm) using a freezing microtome (Leica). Sections were stored at −20 □°C in cryoprotectant solution (ethylene glycol, glycerol, 0.1□M phosphate buffer pH 7.4, 1:1:2 by volume) until used for immunohistochemistry and CARS imaging.

### Immunohistochemistry and BODIPY staining

Free-floating sections were washed three times in PBS, followed by 1 hour blocking in PBS with 10% donkey serum. Sections were incubated in PBS with 10% donkey serum and primary antibodies for 48 hours at 4□°C: rabbit anti-TMEM119 (1:400, Abcam, ab209064), rat anti-CD68 (1:200, Bio-Rad, MCA1957GA), rabbit anti-Iba1 (1:1000, Wako, 019-19741), guinea pig anti-Perilipin 3 (1:200, Progen, G37). After the primary antibody incubation, sections were washed three times in PBS and incubated in PBS with 10% donkey serum and secondary antibodies for 3 hours at room temperature (RT): donkey anti-rabbit Alexa 555, donkey anti-rabbit Alexa 647, donkey anti-rabbit Alexa 405 (all 1:500, Invitrogen), donkey anti-rat Cy5, donkey anti-guinea pig Alexa 488 (all 1:500, Jackson Immuno Research). Sections were washed once in PBS and incubated in PBS with BODIPY^TM^ 493/503 (1:1000 from 1 mg/ml stock solution in DMSO; ThermoFisher) to stain lipid droplets and Hoechst 33342 (1:2000, ThermoFisher) for nuclear counterstaining for 15 min at RT. Sections were mounted on microscope slides and embedded with Vectashield (H-1000, Vector Laboratories). Note that for successful lipid droplet staining, antigen retrieval steps and treatment with detergents have to be avoided, and sections should be embedded while still wet.

### Quantitative analysis of immunohistological data

For quantification of BODIPY^+^ microglia, of CD68^hi^ microglia, of BODIPY^+^ lipid droplet numbers and of lipid droplet size, analyses were performed blinded on coded slides. Every tenth section (400□μm interval) of one hemisphere was selected from each animal and processed for immunohistochemistry. Six randomly selected visual fields per animal were photographed using a confocal scanning laser microscope (LSM 700, Zeiss) with LSM software (ZEN 2011). To analyze the percentage of lipid droplet-containing microglia, numbers of total TMEM119^+^ or Iba1^+^ cells and of TMEM119^+^BODIPY^+^ or Iba1^+^BODIPY^+^ cells were counted and the percentage of BODIPY^+^ microglia was calculated. To assess the percentage of CD68^hi^ cells, BODIPY^-^Iba1^+^ and BODIPY^+^Iba1^+^ cells with clearly visible CD68 immunoreactive particles were counted and normalized to total BODIPY^-^Iba1^+^ and BODIPY^+^Iba1^+^ cells

To assess the numbers of lipid droplets per dentate gyrus area, BODIPY^+^ lipid droplets from six randomly selected visual fields (maximum projection of the z-stack across the whole section) were manually counted and normalized to the corresponding dentate gyrus area. To determine the average size of lipid droplets, BODIPY^+^ signal was analyzed using the ‘Analyze Particles’ function of ImageJ 1.45□s (ImageJ website: http://imagej.nih.gov.laneproxy.stanford.edu/ij/) for six randomly selected visual fields (maximum projection of the z-stack across the whole section).

### 3D Reconstruction of Confocal Images

Confocal image stacks (acquired at 63x magnification) of BODIPY^+^ and CD68^+^ microglia were converted to 3D images with the surface-rendering feature of Imaris BitPlane software (version 7.6.1).

### Human post-mortem brain tissue

Human hippocampal tissue sections from autopsy samples of young adult (<35 years, n=3) and elderly (>60 years, n=5) humans with a post-mortem interval <24□h were used. Human post-mortem tissue was obtained from the collection of the Departmet of Neuropathology at Stanford University. The use of these specimen for scientific purposes was in accordance with institutional ethical guidelines. All samples used were obtained from individuals without any neurological or psychiatric diagnoses. After tissue extraction, the brain samples were stored in 10% formalin. All tissue samples were cut at 50 μm on a vibratome (Leica VT1000S) and stored in PBS at 4□°C

### Immunohistochemistry of formalin fixed human brain tissue

Because immunostaining of formalin-fixed human brain samples requires antigen retrieval steps and the use of detergents (see methods for detailed protocol) which leads to the removal of lipids from tissue, we were unable to use the neutral lipid stain BODIPY. Instead we used an antibody against the lipid droplet surface protein Perilipin2 (Plin2) to detect lipid droplets in human postmortem hippocampus of adult and aged cognitively normal individuals. Formalin-fixed human tissue sections were washed three times in Tris-Buffered Saline with 0.05% Tween 20 (TBST), followed by incubation in TBST with 10% donkey serum and primary antibodies for 72 hours at 4□°C: rabbit anti-Iba1 (1:500, Wako, 019-19741), guinea pig anti-Adipophilin (Plin2) (1:200, Fitzgerald, 20R-AP002). After the primary antibody incubation, sections were washed three times in TBST and incubated in TBST with 10% donkey secondary and secondary antibodies for 3 hours at RT: donkey anti-rabbit Alexa 488, donkey anti-guinea pig Alexa 555 (all 1:500, Invitrogen). Sections were mounted with Vectashield (H-1000, Vector Laboratories).

### CARS Imaging

PFA-fixed 40 μm thick brain sections from 3-month and 20-month old male mice (n = 5 mice per group; 4 sections per animal) were stained for rabbit anti-TMEM119 (1:400, Abcam, ab209064) and donkey-anti rabbit Alexa 647 (Jackson Immuno Research) (Immunohistochemistry protocol see above). Microscopy was performed using a Leica SP5 confocal microscope with spectral detection (Leica Microsystems Inc., Germany) and a Leica HCX PL APO CS 63x NA 1.4 oil immersion objective. Alexa 647 was excited at 633 nm and emission detected between 650-700 nm using a hybrid detector (HyD). Transmission images were acquired simultaneously. Label-free coherent anti-Stokes Raman scattering (CARS) microscopy was performed using a commercial setup consisting of an optical parametric oscillator pumped by a picosecond laser source (*pico*Emerald; APE, Germany) integrated into the Leica SP5 microscope. The CARS signal was detected using a 650/210 bandpass emission filter and a non-descanned detector (NDD) in epi-mode. To detect neutral lipids the laser was tuned to 2845 cm^-1^, thus enabling imaging of CH_2_ symmetric stretching vibrations. CARS and fluorescence/transmission images were acquired sequentially. For quantification of CARS signal in microglia, 20 randomly selected microglia in the dentate gyrus per animal were imaged. The percentage of TMEM119^+^ microglia with CARS^+^ vesicles from total TMEM119^+^ microglia was calculated.

### Lipidomics

5-month and 20-month old male wild type mice (n = 4 mice/ group) were perfused and hippocampus and liver were extracted. Alternatively, hippocampal microglia were FACS-sorted from 20-month old mice (see “Microglia Isolation” above). BV2 cells were treated for 18□hours with 5 μg/ml LPS (Lipopolysaccharide from E. coli, Sigma) in DMEM +5% FBS to induce lipid droplet formation. Lipid droplets from liver, whole hippocampus, isolated microglia, and LPS-treated BV2 cells were isolated using the lipid droplet isolation kit from Cell Biolabs according to the manufacturer’s instructions.

Lipid droplets were stored at -70°C until sample preparation and extracted according to a modified Bligh & Dyer protocol^2^ and as reported earlier^7^. Prior extraction, 10 µL of a synthetic lipid standard mastermix (including 15 deuterated lipids) were added to 90 µL of extraction buffer containing lipid droplets.

Extracted lipids were analyzed by flow injection analysis (FIA) shotgun lipidomics using an ekspert™ MicroLC 200 system (eskigent, Singapore) hyphenated to a TripleTOF® 4600 System (AB SCIEX, Darmstadt, Germany) as reported earlier^8^. Each sample was injected twice, one for measurement in positive and one for negative ionization mode. Instrumental controlling and data acquisition was achieved with Analyst® TF Software (v 1.7, AB SCIEX, Darmstadt, Germany). Data were processed with Lipid View™ software (v 1.3 beta, AB SCIEX, Darmstadt, Germany). Lipid identification was based on precursor ion and neutral loss scans specific for proposed lipid species. Internal standard correction for each lipid was carried out by normalization against the appropriate synthetic isotopically labeled lipid standard.

### Microglia Isolation

Primary microglia were isolated as previously described^3^. 18-month-old male wild type mice were perfused with Medium A (HBSS+ 15mM HEPES+0.05% glucose+ 1:500 DnaseI), and hippocampi were dissected (n = 9 mice; 3 pooled hippocampi per sample). Hippocampi were chopped and homogenized using a Dounce homogenizer in 2 ml cold Medium A (HBSS+ 15mM HEPES+0.05% glucose+ 1:500 DnaseI), filtered through 100 μm cell strainer, rinsed with 5 ml Medium A and centrifuged at 340 g for 5 min. For myelin removal, the precipitate was resuspended in 30% standard isotonic percoll (30% Percoll in PBS, diluted with Medium A) and centrifuged at 900 g for 20 min. Precipitated cells were washed with HBSS and resuspended in FACS buffer (PBS+1% BSA+2mM EDTA). The samples were stained with 1:300 CD11b-PE and 1:300 CD45-APC for 30 min at RT, centrifuged at 400 g for 5 min, resuspended in PBS with BODIPY 493/503 (1:2000 from 1 mg/ml stock solution in DMSO; ThermoFisher) and incubated for 10 min at 37°C. Cells were washed two times with FACS buffer and resuspended in FACS buffer with DNAse1 and 5 μl/ml RNAse Inhibitor (Clontech). Dead cells were excluded by staining with Sytox blue dead cell stain (1:10,000, Invitrogen). Cells were isolated using an ARIA 3.1 (BD Biosciences) with FACSDiva software (BD Biosciences), were sorted into RLT lysis buffer (QIAGEN) with 1% 2-Mercaptoethanol and frozen at -80°C.

### RNA isolation and library preparation

Frozen cells were thawed to room temperature and total RNA was isolated from the cell pellets using the RNeasy Plus Micro kit (Qiagen, 74034). RNA quantities and RNA quality was assessed using the Agilent 2100 Bioanalyzer (Agilent Technologies). All samples passed a quality control threshold (RIN ≥ 9.0) to proceed to library preparations and RNAseq. Total mRNA was transcribed into full length cDNA using the SMART-Seq v4 Ultra Low Input RNA kit from Clontech according to the manufacturer’s instructions. Samples were validated using the Agilent 2100 Bioanalyzer and Agilent High Sensitivity DNA kit. 150 pg of full length cDNA was processed with the Nextera XT DNA library preparation kit from Illumina according to the manufacturer’s protocol. Library quality was verified with the Agilent 2100 Bioanalyzer and the Agilent High Sensitivity DNA kit. Sequencing of microglia isolated from aged wild type mice was carried out with Illumina HiSeq 2000/2500, paired end, 2x 100 bp depth sequencer, and microglia from 10-month-old GRN^-/-^ mice and from 3-month-old LPS-treated mice were sequenced with the Illumina Novogene 6000, paired end, 2x 100 bp. The quality of fastq files was assessed using FASTQC (v 0.11.4). Reads were mapped to mouse mm9 reference genome using STAR (v 2.5.1b). Raw read counts were generated with STAR using the GeneCounts function.

### RNA-seq differential expression

Differential expression in RNA-Seq was analyzed using the R DESeq2 package (Love et al., 2014). Read counts were used as input and normalized using built-in algorithms in DESeq2. Pairwise comparisons among the two groups (BODIPY^lo^ and BODIPY^hi^ microglia) were done on all genes and 12129 genes with calculable fold changes (FC) and false discovery rates (fdr) were used for further analysis. False discovery rate was estimated using Benjamini and Hochberg approach (Benjamini and Hochberg, 1995). R was used for RNA-Seq data visualization and Ingenuity Pathway Analysis (IPA) and Enrichr were used to analyze pathways and upstream regulators.

For comparisons of transcriptome changes in LAM with published datasets (Fig 2g and Supplementary Figure 3), we selected the following published RNA-seq datasets of microglia in aging and neurodegeneration (modified after Bohlen et al. 2017): 24-month versus 4-month old wild type mice for Aging (Holtman et al. 2015); APP+ versus APP-for AD (Wang et al., 2015); SOD^G93A^ endstage versus non-transgenic day 130 for ALS (Chiu et al., 2013); DAM versus homeostatic microglia (Keren-Shaul et al. 2017), MGnD versus homeostatic microglia (Kraseman et al. 2017), and microglia clusters published by Li et al. (2019) and Hammond and colleagues (2019). For gene set comparisons, we generated lists for the published datasets by using a 4-fold change cutoff (modified after Bohlen et al. 2017) and compared these lists with the top 100 up- and downregulated genes in LAM.

### BV2 cell culture

Cells from the murine microglial cell line BV2 were originally obtained from Banca Biologica e Cell Factory, IRCCS Azienda Ospedaliera Universitaria San Martino, Genua, Italy. Cells were maintained in Dulbecco’s Modified Eagle’s Medium (DMEM, Life Technologies) supplemented with 10% fetal bovine serum (FBS) and antibiotics (penicillin 100□U□ml^−1^, streptomycin 100□U□ml^−1^, HVD Life Sciences) (Pen/Strep) under standard culture conditions (95% relative humidity with 5% CO_2_ at 37°C). Adherent cells were split using 1X TrypLE (GIBCO).

### LPS and Triacsin C treatment

To induce lipid droplet formation, subconfluent BV2 cells were treated for 18□hours with 5 μg/ml LPS (Lipopolysaccharide from *E. coli*, Sigma) in DMEM +5% FBS. Controls received vehicle solution (PBS) only. To inhibit lipid droplet formation, BV2 cells were pre-treated with 1 μM Triacsin C (Cayman Chemical) or vehicle (saline) in DMEM +5% FBS. 30 min after adding Triacsin C, LPS or LPS-vehicle solution were added and cells were co-treated with Triacsin C and LPS for 18 hours.

### Plasma collection

Blood from young (2-month-old) and aged (18-month-old) wild type mice was collected with ethylene diamine tetra acetate (EDTA) as anticoagulant via intracardial bleed at time of death. EDTA plasma was obtained from freshly collected blood by centrifugation (1000 g for 10 min at 4 °C), aliquoted, and stored at -80 °C until use. For BV2 experiments, thawed plasma was dialyzed in PBS to remove EDTA, and then delipidized using a Lipid Removal Adsorbent (LRA, Sigma). Briefly, plasma was mixed with LRA (40mg/mL) for 60 minutes at RT, centrifuged at 2200 x g for 2 minutes, and the supernatant was collected. Dialyzed and delipidized plasma was diluted to working concentration (5%) in DMEM +5% FBS and incubated for 20 min at room temperature to allow to clot. The solution was filtered through a PES 0.22um filter unit and used for cell culture assays.

### Plasma treatment

BV2 cells were treated for 18□hours with 5% plasma (preparation see *Plasma collection*) in DMEM +5% FBS. Controls received vehicle solution (PBS) only.

### BODIPY *in vitro* staining

BV2 cells were seeded at 5x10^4^ cells on poly-L-lysine coated glass coverslips in DMEM +5% FBS. Following specific treatments, cells were fixed in 4% PFA for 30 min, washed 3x in PBS and incubated in PBS with BODIPY^TM^ 493/503 (1:1000 from 1mg/ml stock solution in DMSO; ThermoFisher) and Hoechst 33342 (1:2000, ThermoFisher) for 10 min at RT. Sections were washed twice in PBS and mounted on microscope slides with Vectashield (H-1000, Vector Laboratories). Four randomly selected visual fields per coverslip were photographed (40x magnification) using a confocal scanning laser microscope (LSM 700, Zeiss) with LSM software (ZEN 2011). To analyze the percentage of lipid droplet-containing BV2 cells, numbers of total Hoechst^+^ cells and of Hoechst^+^ cells with BODIPY^+^ lipid droplets were counted and the percentage of BODIPY^+^ BV2 cells was calculated.

### *In vitro* phagocytosis assay

For *in vitro* phagocytosis assays, BV2 cells were split into 96 well plates at 1000 cells per well in DMEM +5% FBS and treated with LPS, Triacsin C and vehicle solutions, or with 5% plasma and vehicle, for 18 hours. Following specific treatments, 5 ng pHRodo Red Zymosan™ Bioparticles (Thermo Fisher Scientific P35364) in 100 μl DMEM +5% FBS were added per well. Four phase contrast and red fluorescent images per well were acquired every 2 hours for 16 hours using the Incucyte S3 live cell analysis system (Essen Bioscience). For each time point, phagocytosis was calculated by normalizing red fluorescent area to phase confluence.

### Organotypic brain slices and *in situ* phagocytosis assay

12-month-old male wild type mice (n = 3 mice) and 9-month-old male GRN^-/-^ mice were decapitated and dissected brains were immediately put in pre-cooled culture medium with serum (65% MEM (Sigma); 10% horse serum; 25% HBSS; 6.5 mg/ml glucose; 2mM Glutamine; 1% Pen/Strep).

The entire procedure was done on ice with pre-cooled solutions until culturing. Coronal sections were prepared with a vibratome (Leica VT1000S) at 250 μm thickness and then transferred to insert wells (Millicell Cell Culture Insert, 30 mm, Millipore) on a 6-well plate with medium.

Sections were incubated for 1 hour in the incubator (37°C, 5% CO_2_). pHrodo™ Red Zymosan Bioparticles (Thermo Fisher Scientific) were opsonized (3 washes in PBS, followed by incubation in 50% FBS in PBS for 45 min at 37°C and three washes in PBS) and added at 0.5 mg/ml to cover the whole sections (about 150 μl each). The plate was incubated for 4 hours (37°C, 5% CO2). After washes with PBS, sections were fixed with 4% PFA for 30 min at RT.

Immunohistochemistry for Iba1 and BODIPY staining was performed as described above. To measure phagocytosis in BODIPY^-^ and BODIPY^+^ microglia, four randomly selected visual fields per section (three sections per mouse) were photographed using a confocal scanning laser microscope (LSM 700, Zeiss) with LSM software (ZEN 2011). The numbers of BODIPY^-^Iba1^+^ and BODIPY^+^Iba1^+^ cells were quantified, and the percentage of Zymosan containing cells was calculated.

### *In vivo* phagocytosis assay

CypHer5E and AlexaFluor555 double-labelled myelin (25 mg/ml in PBS) was injected into the hippocampus of 20-month old male mice using a stereotaxic apparatus (Kopf instruments). Mice were anaesthetized using isofluorane, their skulls were exposed and a hole was drilled at the injection site using aseptic technique. One microliter of the myelin solution was injected at ±0.7 mm lateral, -1.7 mm anterior–posterior, and −2.04 mm dorso-ventral relative to the intersection of the coronal and sagittal suture (bregma) at a rate of 200 nl/min. The needle was left in place for an additional 3 min to allow for diffusion, then slowly withdrawn. Mice received post-surgical buprenorphine and baytril for pain and infection prevention, respectively. After 48 h, mice were anaesthetized and transcardially perfused with 4% PFA. The entire injection site was sectioned (coronal, 40 μ thick) and stained for Iba1 and BODIPY as described above. 5-8 sections were quantified to assess myelin uptake of BODIPY^+^/Iba1^+^ cells and BODIPY^-^/Iba1^+^ cells.

### ROS assays

To assess ROS generation in primary microglia, cell homogenates from 3- and 20-month-old male wild type mice and from 9-month old male GRN^-/-^ mice were prepared and antibody staining was performed as described above (see *Microglia Isolation*). Cell homogenates were incubated in FACS buffer with CellROX^TM^ Deep Red (1:500, Invitrogen) for 30 min at 37°C, washed twice in FACS buffer, and CellROX^TM^ Deep Red Intensity was analyzed on ARIA 3.1 (BD Biosciences). To measure ROS in BV2 cells, cells were split into 24 well plates at 5x10^4^ cells per well in DMEM +5% FBS and treated with LPS, Triacsin C and vehicle solutions for 18 hours. Next, cells were incubated in DMEM+5%FBS with CellROX^TM^ Orange (1:500, Invitrogen) for 30 min at 37°C, washed twice in PBS, and CellROX^TM^ Orange Intensity was examined by fluorescence microscopy (Keyence Corp., Osaka, Japan). Following microscopic analysis, cells were detached using TripLE, transferred to FACS tubes and CellROX^TM^ Orange intensity was analyzed on BD AccuriC6 flow cytometer.

### iodo-BODIPY irradiation assay

BV2 cells were treated for 18 hours with 5 μg/ml LPS in DMEM with 5% FBS to induce lipid droplet formation. Controls re^L^ceived vehicle solution (PBS) only. Cells were washed twice in PBS and incubated in 3.8 μM iodo-BODIPY (see Figure 6a; iodo-BODIPY was kindly provided by Prof. Carolyn Bertozzi, Chemistry department, Stanford University) in PBS for 30 min in the incubator (5%CO_2_, 37°C). Cells were washed twice in PBS and then irradiated under visible light (green LED; 9W, 150mA, 2000 lux) for 3 hours at RT. Non-irradiated cells were kept at RT for 3 hours without irradiation.

### CalceinAM live cell staining

To assess cell viability after photoirradiation, irradiated and non-irradiated BV2 cells were stained with 1μM Calcein AM (Invitrogen) in PBS for 15 min at 37°C. Cells were then examined by fluorescence microscopy (Keyence Corp., Osaka, Japan) and the percentage of green, Calcein^+^ BV2 cells (live cells) of total methyl-di-iodo BODIPY^+^ cells (red, lipid droplet containing cells) was calculated.

### CRISPR-Cas9 Screen

The 10-sgRNA-per-gene CRISPR/Cas9 deletion library was synthesized, cloned and infected into Cas9-expressing BV2 cells as previously described^4^. Briefly, about 30 million BV2 cells stably expressing EF1-alpha-Cas9-Blast were infected with the 10 guide/gene sgRNA sub-library (see Supplementary Table 2 for gene list) at an MOI<1. Infected cells underwent puromycin selection (1.5μg/ml) for 7 days after which point puromycin was removed and cells were resuspended in normal growth medium (DMEM+10%FBS) without puromycin. After selection, sgRNA infection was confirmed by flow cytometry, which indicated >90% of cells expressed the mCherry reporter. Cells were cultured at 1000x coverage (about 1000 cells containing each sgRNA) throughout the screen. BV2 cells were treated for 18 hours with 5 μg/ml LPS in DMEM with 5% FBS to induce lipid droplet formation, and the photoirradiation assay was performed as described above (see *Methyl-di-iodo BODIPY irradiation assay*). Non-irradiated, LPS-treated BV2 cells were used as controls. After irradiation, cells were washed twice in medium and put back in the incubator. LPS-treatment and photoirradiation were performed three times, with 24 hours recovery time between each irradition and the next LPS treatment. At the end of the screen genomic DNA was extracted for all experimental conditions using a QIAGEN Blood Midi Kit. Deep sequencing of sgRNA sequences on an Illumina Nextseq was used to monitor library composition. Guide composition was analyzed and compared to the plasmid library between conditions using castle version 1.0^5^ available at https://bitbucket.org/dmorgens/castle. Genes with FDR<50% were considered as significant hits.

### Generation of single CRISPR-Cas9 knockout BV2 cells

Lentivirus production and infection was performed as previously described^4^. Briefly, HEK293T cells were transfected with packaging plasmids and sgRNA-containing plasmids. Supernatant was harvested at 48 and 72 hours and concentrated with Lenti-X solution (Clontech). BV2 cells stably expressing Cas9 under blasticidin (1 μg/ml) were infected with lentivirus containing sgRNA plasmids under puromycin selection for 24 hours. Puromycin selection was started 48 hours after infection and maintained for 7 days. *GRN* knockout BV2 cells were sub-cloned to maintain a monoclonal knockout population. *SLC33A1* and *VPS35* single-knockout cell lines were assayed as polyclonal populations.

### Cytokine assay

Primary microglia from hippocampi of 3-month (young) and 20-month-old (aged) male wild type mice and from 10-month-old male GRN^-/-^ mice were isolated as described above (see *Microglia Isolation*, hippocampi from 3 mice were pooled per age group). CD11b^+^CD45^lo^ primary microglia from young mice, and CD11b^+^CD45^lo^ primary microglia sorted for BODIPY^lo^ and BODIPY^hi^ cells from aged mice were sorted into 5% FBS-containing microglial culture medium (DMEM/F12, 1% Pen/Strep; 2 mM glutamine; 5 ug/mL N-acetyl cysteine; 5 ug/mL insulin; 100 ug/mL apo-transferrin; 100 ng/mL sodium selenite; 2 ng/mL human TGF-2; 100 ng/mL murine IL-34; 1.5 ug/mL ovine wool cholesterol; 10 ug/mL heparin sulfate)^6^. Cells were seeded into 96 well plates at 5000 cells per well in 100 μ microglia culture medium with 5%FBS and incubated for 30 min in the incubator (37°C, 5% CO_2_). Next, cells were treated with 100 ng/ml LPS or PBS for 8 hours, and supernatant was collected and secreted signaling proteins were measured in culture supernatants using the ‘Mouse Cytokine / Chemokine Array 31-Plex’ from Eve Technologies (Canada).

### NAD/NADH assay

Primary microglia from 3-month (young) and 20-month-old (aged) wild type mice were isolated as described above (see *Microglia Isolation;* hippocampi from 3 mice were pooled per group), and CD11b^+^CD45^lo^ primary microglia from young mice, and CD11b^+^CD45^lo^ primary microglia sorted for BODIPY^lo^ and BODIPY^hi^ cells from aged mice were sorted into 5% FBS-containing microglial culture medium (see *Cytokine assay*). Cells were seeded into 96 well white-walled 52μl microglial culture medium with 5%FBS and tissue culture plates at 5000 cells per well in 50 μl microglial culture medium with 5%FBS and incubated for 30 min in the incubator (37°C, 5% CO_2_). The NAD/NADH-Glo^TM^ assay (Promega) was performed according to manufacturer’s instructions. Cell lysates were incubated for 2 hours at RT and luminescence was recorded using a luminometer (Lmax, Molecular Devices).

### Statistical analysis

Data collection was randomized for all experiments. Experimenters were blinded for imaging and data analysis. Statistical analyses were performed using the GraphPad Prism 5.0 software (GraphPad Software), and R DESeq2 package was used for RNA-Seq analysis. Data were tested for normality using the Kolmogorov–Smirnov or the Shapiro–Wilk test, and equality of variance was confirmed using the F-test. Means between two groups were compared by the two-tailed unpaired Student’s *t*-test or, in case of non-gaussian distribution, by using the two-tailed Mann– Whitney *U*-test. Data from multiple groups were analyzed by one-way ANOVA and two-way ANOVA, followed by Tukey’s *post hoc* tests. Detailed information on sample size, numbers of replicates, and statistical test used for each experiment is provided in the figure legends.

### Data availability

Raw data are available from the corresponding author upon request.

