## Supplemental Figures 1-7 for "Lipid droplet accumulating microglia represent a dysfunctional and pro-inflammatory state in the aging brain"

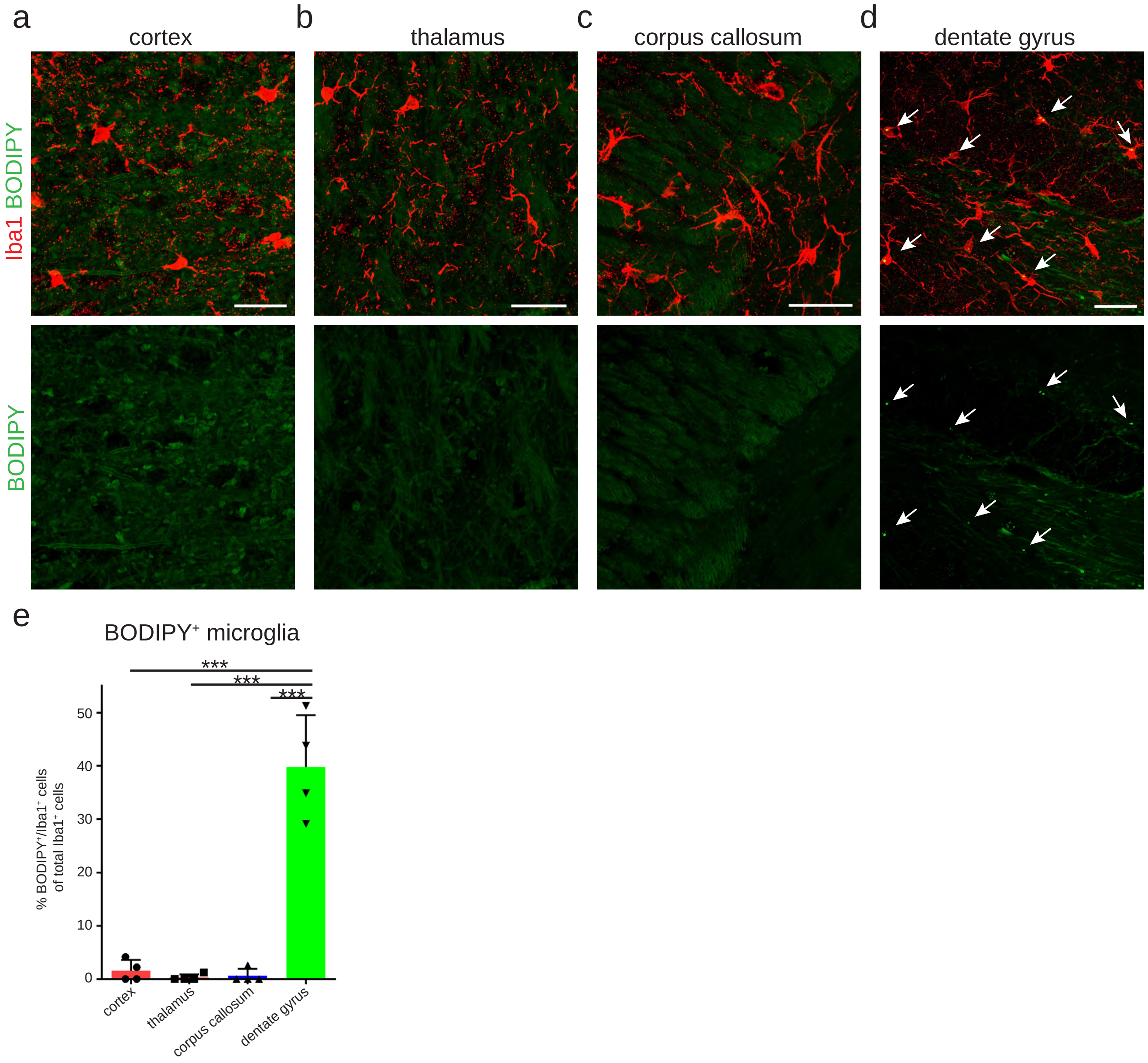
a
cortex
b
thalamus
c
corpus callosum
d
dentate gyrus
Iba1 BODIPY
BODIPY
e
BODIPY⁺ microglia
***
***
***
% BODIPY⁺/Iba1⁺ cells of total Iba1⁺ cells
50
40
30
20
10
0
cortex
thalamus
corpus callosum
dentate gyrus

### Supplementary Figure 1

**Supplementary Fig. 1 Lipid droplet accumulating microglia are abundant in the hippocampus but rare in other brain regions of aged mice.** *Related to Figure 1*
**a-d**, Representative confocal images of the cortex (**a**), thalamus (**b**), corpus callosum (**c**) and hippocampal dentate gyrus (**d**) from 20-month old male mice stained for BODIPY$^+$ (lipid droplets) and Iba1$^+$ (microglia). Scale bar: 20 µm. Arrows point towards BODIPY$^+$ lipid droplets. e, Quantification of BODIPY$^+$/Iba1$^+$ cells. n = 4 mice per group. One-way ANOVA followed by Tukey's post hoc test. Error bars represent mean ± SD. ***P< 0.001.

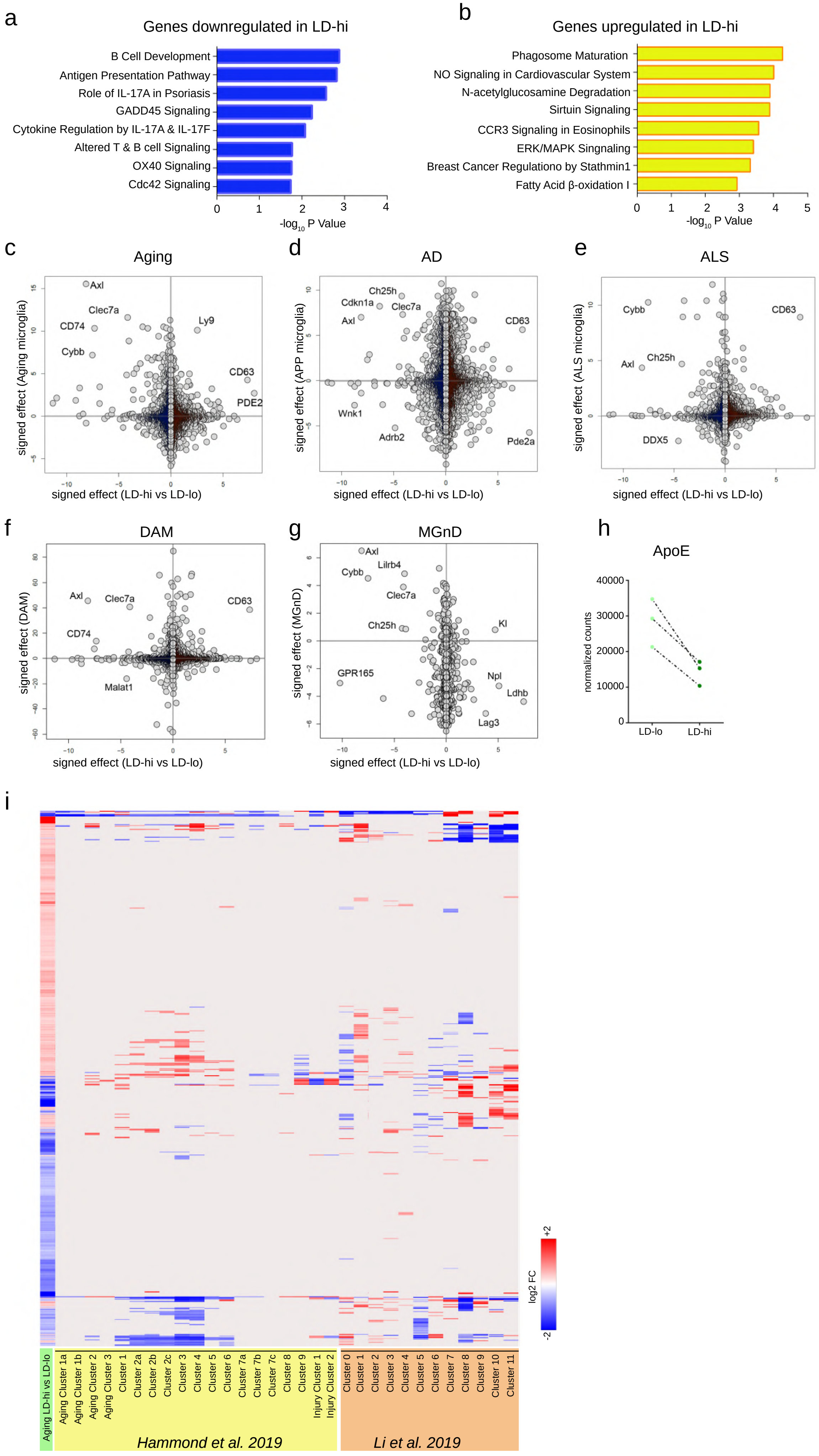

a
Genes downregulated in LD-hi
B Cell Development
Antigen Presentation Pathway
Role of IL-17A in Psoriasis
GADD45 Signaling
Cytokine Regulation by IL-17A & IL-17F
Altered T & B cell Signaling
OX40 Signaling
Cdc42 Signaling
-log10 P Value
b
Genes upregulated in LD-hi
Phagosome Maturation
NO Signaling in Cardiovascular System
N-acetylglucosamine Degradation
Sirtuin Signaling
CCR3 Signaling in Eosinophils
ERK/MAPK Singnaling
Breast Cancer Regulationo by Stathmin1
Fatty Acid β-oxidation I
-log10 P Value
c
Aging
signed effect (Aging microglia)
signed effect (LD-hi vs LD-lo)
d
AD
signed effect (APP microglia)
signed effect (LD-hi vs LD-lo)
e
ALS
signed effect (ALS microglia)
signed effect (LD-hi vs LD-lo)
f
DAM
signed effect (DAM)
signed effect (LD-hi vs LD-lo)
g
MGnD
signed effect (MGnD)
signed effect (LD-hi vs LD-lo)
h
ApoE
normalized counts
LD-lo
LD-hi
i
log2 FC
Aging LD-hi vs LD-lo
Hammond et al. 2019
Li et al. 2019

**Supplementary Fig. 2 LAM have a unique transcriptional signature that minimally overlaps with published gene expression profiles of microglia in aging and neurodegeneration.** *Related to Figure 2*

**a**,**b**, IPA pathway analysis of genes that are significantly upregulated (a) or downregulated (b) in LD-hi microglia in aging. Analysis based on top 100 down- and up-regulated genes. **c**-**g**, Expression plots comparing RNA-Seq data of LAM (see Fig. 2) with published RNA-Seq data of microglia in aging (**c**), AD (**d**), ALS (**e**), disease-associated microglia (DAM) (**f**) and neurodegenerative microglia (MGnD) (**g**). Data are expressed as signed fdr, i.e the product of log2 FC and log10 fdr. **h**, Paired dot plot showing FPKM values of LD-lo and LD-hig microglia for ApoE (P= 0.423). Dotted lines connect LD-lo and LD-hi microglia sorted from the same samples. **i**, Heatmap showing expression changes of LAM genes (genes differentially expressed in LD-hi microglia in aging) in LD-hi microglia from $GRN^{-/-}$ mice, from LPS treated mice, and in microglia clusters revealed by Li et al. (2019) and Hammond et al. (2019)[14,15]. LD, lipid droplet.

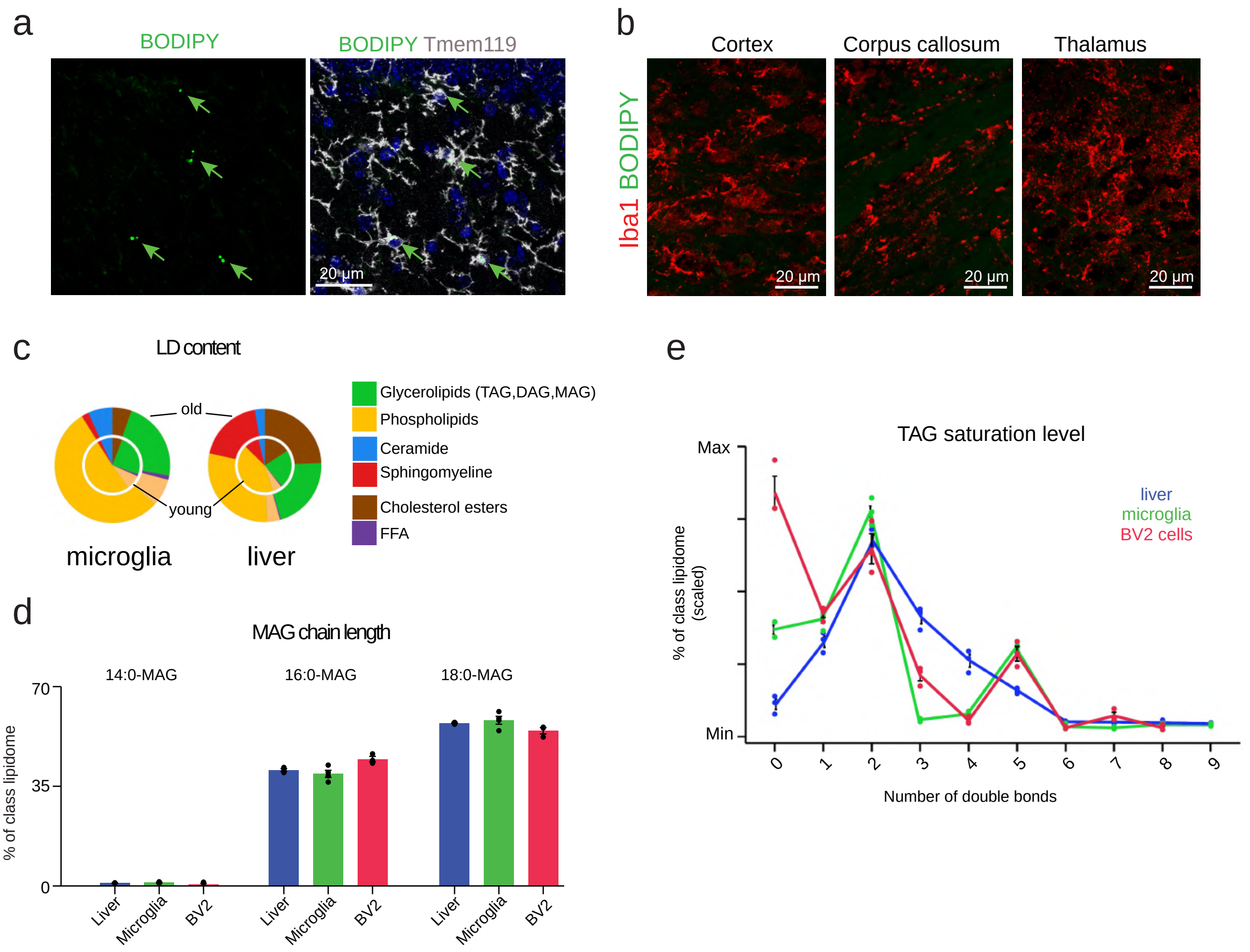
a
BODIPY
BODIPY Tmem119
20 µm
b
Cortex
Corpus callosum
Thalamus
Iba1 BODIPY
20 µm
20 µm
20 µm
c
LD content
old
young
microglia
liver
Glycerolipids (TAG,DAG,MAG)
Phospholipids
Ceramide
Sphingomyeline
Cholesterol esters
FFA
d
MAG chain length
14:0-MAG
16:0-MAG
18:0-MAG
70
35
0
% of class lipidome
Liver
Microglia
BV2
Liver
Microglia
BV2
Liver
Microglia
BV2
e
TAG saturation level
Max
Min
% of class lipidome (scaled)
liver
microglia
BV2 cells
0
1
2
3
4
5
6
7
8
9
Number of double bonds

**Supplementary Fig. 3 LPS treatment induces lipid droplet formation in microglia and in BV2 cells.** *Related to Figure 3*

**a**,**b**, 3-month-old male mice were given intraperitoneal (i.p.) injections of LPS (1 mg/kg BW) for four days. Representative confocal images of BODIPY$^{+}$ and Tmem119$^{+}$ in the hippocampus (**a**) and of BODIPY and Iba1 staining in the cortex, corpus callosum, and thalamus (**b**). **c-e**, Lipidome profiling of lipid droplets from LPS-treated BV2 cells, primary microglia, and liver tissue. **c**, Pie charts showing that the lipid composition of lipid droplets from young and aged microglia is highly similar, but differs between young and aged liver tissue. **d**,**e**, Distribution of MAG chain lengths (**d**) and TAG saturation levels (**e**) of lipid droplets isolated from LPS-treated BV2 cells and from microglia and liver tissue from aged mice. young= 5-month-old male mice, old= 20-month-old male mice; n = 4 mice per group.

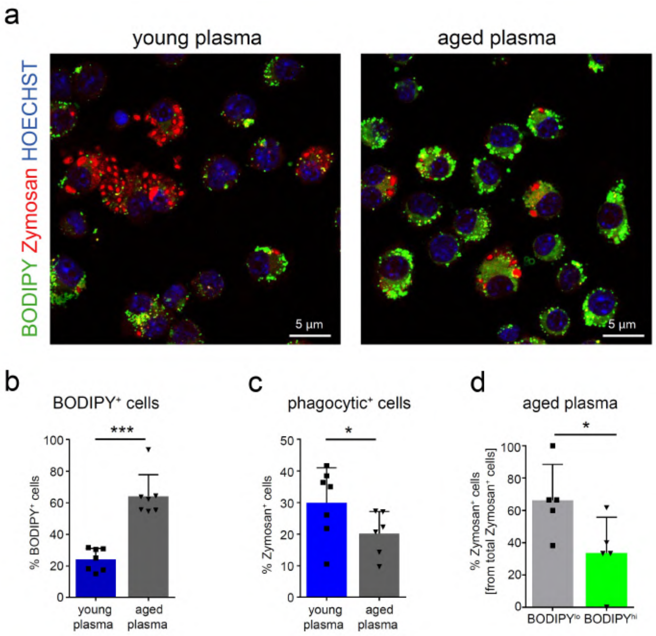
a
young plasma
aged plasma
BODIPY Zymosan HOECHST
5 µm
5 µm
b
BODIPY⁺ cells
% BODIPY⁺ cells
***
young plasma
aged plasma
c
phagocytic⁺ cells
% Zymosan⁺ cells
*
young plasma
aged plasma
d
aged plasma
% Zymosan⁺ cells [from total Zymosan⁺ cells]
*
BODIPY^lo
BODIPY^hi

**Supplementary Fig. 4 Aged plasma induces lipid droplet formation in BV2 cells.** *Related to Figure 4*

**a**, Representative micrographs of $BODIPY^+$ staining and of phagocytosis of of pHrodo red Zymosan in BV2 cells treated with 5% plasma from young (3-months) and aged (18-months) mice for 12 hours. **b**, Quantification of $BODIPY^+$ staining in BV2 cells treated with young and aged plasma. **c**,**d,** Quantification of Zymosan uptake in BV2 cells treated with young and aged plasma (**c**), and in aged plasma treated BODIPY-low and BODIPY-high cells (**d**). Statistical tests: unpaired Student's t-test. Error bars represent mean ± SD. *$P< 0.05$, ***$P< 0.001$.

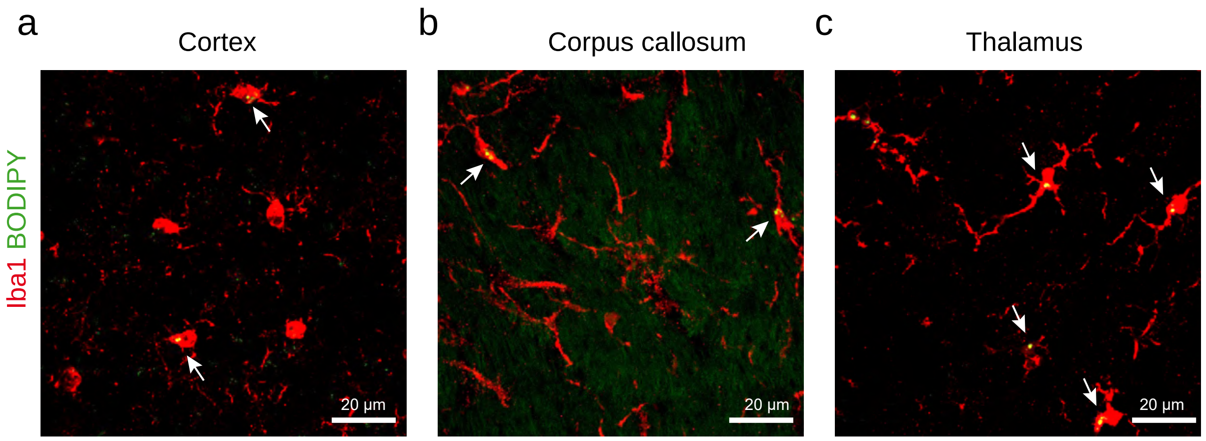
a
Cortex
b
Corpus callosum
c
Thalamus
Iba1 BODIPY
20 μm
20 μm
20 μm

**Supplementary Fig. 5 Lipid droplet containing microglia in the cortex, corpus callosum, and thalamus of GRN$^{-/-}$ mice.** *Related to Figure 7*

**a**-**c**, Representative confocal images of BODIPY$^{+}$ (lipid droplets) and Iba1$^{+}$ (microglia) in the cortex (**a**), corpus callosum (**b**), (**c**) and thalamus from 9-month-old male GRN$^{-/-}$ mice. BODIPY$^{+}$/Iba1$^{+}$ cells were frequently found in the thalamus and were detected to a lesser extent in cortex and corpus callosum.

**a**

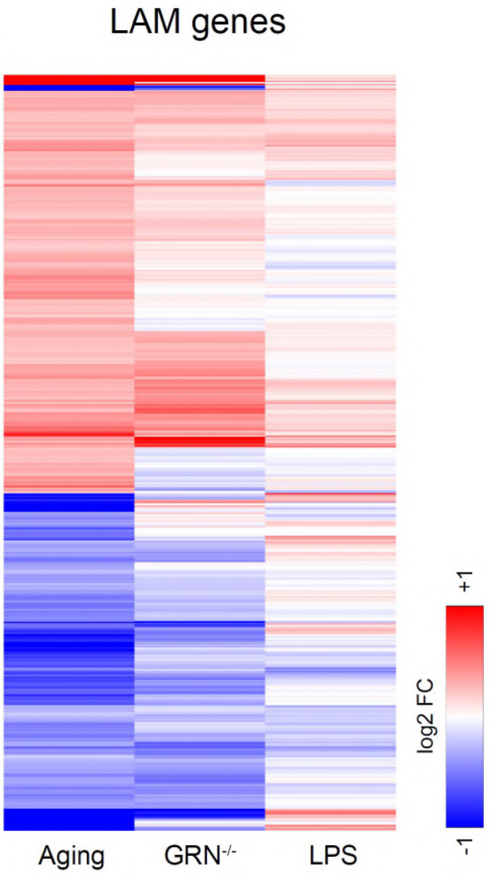

**Supplementary Fig. 6 Expression changes of LAM genes in lipid droplet-rich microglia from normal aging, GRN$^{-/-}$ and LPS-treated mice.** *Related to Figures 3 and 7*

**a**, Heatmap showing expression changes of LAM genes (genes differentially expressed in LD-hi microglia in aging; 692 genes) in LD-hi microglia from GRN$^{-/-}$ mice and from LPS treated mice.

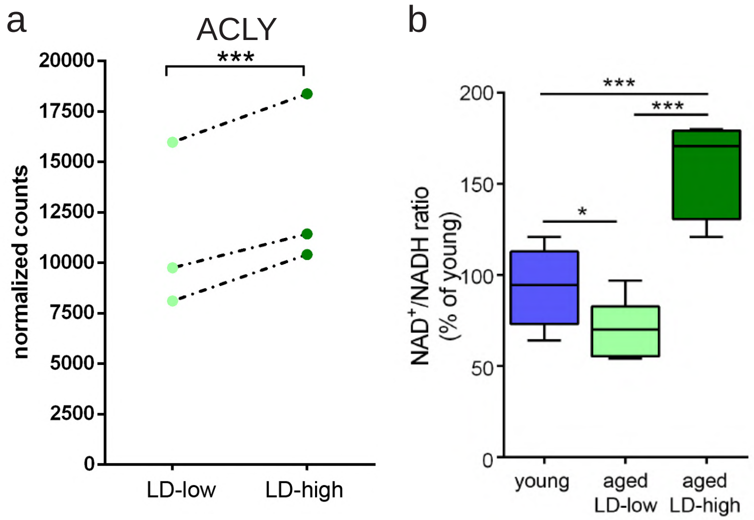
a
ACLY
***
normalized counts
20000
17500
15000
12500
10000
7500
5000
2500
0
LD-low
LD-high
b
***
***
*
200
150
100
50
0
NAD⁺/NADH ratio
(% of young)
young
aged
LD-low
aged
LD-high

**Supplementary Fig. 7 LAM show signs of metabolic alterations.** *Related to Figure 2*
**a**, Paired dot plot showing FPKM values of LD-lo and LD-hi microglia for ACLY (data obtained from RNA-Seq analysis, see Figure 2). Dotted lines connect LD-lo and LD-hi microglia sorted from the same samples. **b**, NAD colorimetric assay showing the $NAD^+$/NADH ratio of primary hippocampal microglia from 3-month old mice (young) and of LD-lo and LD-hi primary microglia from 20-month old male mice. Experiments were performed two times in technical triplicates. n=3 mice per group per experiment. Statistical tests: one-way ANOVA (b) followed by Tukey's post hoc test. Horizontal lines in the box plots indicate medians, box limits indicate first and third quantiles, and vertical whisker lines indicate minimum and maximum values. *P< 0.05, ***P< 0.001.
